# Silencing of *Mu* elements in maize involves distinct populations of small RNAs and distinct patterns of DNA methylation

**DOI:** 10.1101/2020.01.09.900282

**Authors:** Diane Burgess, Hong Li, Meixia Zhao, Sang Yeol Kim, Damon Lisch

## Abstract

Epigenetic changes involve changes in gene expression that can be heritably transmitted to daughter cells in the absence of changes in DNA sequence. Epigenetics has been implicated in phenomena as diverse as development, stress response and carcinogenesis. A significant challenge facing those interested in investigating epigenetic phenomena is determining causal relationships between DNA methylation, specific classes of small RNAs and associated changes in gene expression. Because they are the primary targets of epigenetic silencing in plants and, when active, are often targeted for *de novo* silencing, transposable elements (TEs) represent a valuable source of information about these relationships. We use a naturally occurring system in which a single TE can be heritably silenced by a single derivative of that TE. By using this system it is possible to unravel causal relationships between different size classes of small RNAs, patterns of DNA methylation and heritable silencing. Here, we show that the long terminal inverted repeats (TIRs) within *Zea mays MuDR* transposons are targeted by distinct classes of small RNAs during epigenetic silencing that are dependent on distinct silencing pathways. Further, these small RNAs target distinct regions of the TIRs, resulting in different patterns of cytosine methylation with different functional consequences with respect to epigenetic silencing and heritability of that silencing.

**Summary:** Transposable elements (TEs) are a ubiquitous feature of plant genomes. Because of the threat they post to genome integrity, most TEs are epigenetically silenced. However, even closely related plant species often have dramatically different populations of TEs, suggesting periodic rounds of activity and silencing. Here we show that the process of *de novo* methylation of an active element in maize involves two distinct pathways, one of which is directly implicated in causing epigenetic silencing and one of which is the result of that silencing.

## Introduction

Plant genomes are host to large numbers of potentially deleterious endogenous mutagens known as transposable elements (TEs). Due to the activity of a sophisticated regulatory system, the vast majority of TEs are epigenetically silenced (Slotkin and Martienssen 2007; Lisch 2009; Law and Jacobsen 2010; Bucher *et al*. 2012; Sigman and Slotkin 2016). This silencing is associated with DNA methylation and modification of histones, and can be propagated over many generations (Lisch 2009; Saze and Kakutani 2011).

Because most transposons are silenced most of the time, much of what we know about TE silencing involves maintenance, rather than initiation of silencing. However, recent work suggests that aberrant RNAs can trigger silencing of otherwise active TEs via a pathway that involves the production of *trans*-acting 21-22 nt small RNAs via the activity components of both the Post Transcriptional Gene Silencing (PTGS) and the Transcriptional Gene Silencing (TGS) pathways (Li *et al*. 2010; Mari-Ordonez *et al*. 2013; Nuthikattu *et al*. 2013; Fultz *et al*. 2015; Cuerda-Gil and Slotkin 2016). The available data suggests that Pol II transcripts from active TEs are recognized by small RNAs that then act as triggers for RDR6/SGS3-mediated production of dsRNAs using RNA directed RNA polymerase 6 (RDR6) in conjunction with Suppressor of Gene Silencing 3 (SGS3). The resulting double-stranded RNA is then processed into 21-22 nt *trans*-acting small RNAs by Dicer like 2 (DCL2) or Dicer like 4 (DCL4). These small RNAs are then incorporated into a complex that includes Argonaute 6 (AGO6), which is then competent to trigger *de novo* and heritable TGS using Pol V transcript arising from the active elements as a scaffold (Fultz *et al*. 2015; McCue *et al*. 2015; Cuerda-Gil and Slotkin 2016; Fultz and Slotkin 2017). The initial triggers for silencing are not always well understood, but it appears that they may be small RNAs derived from the Pol II transcript itself, or from an unlinked aberrant version of the TE (Slotkin *et al*. 2005; Mari-Ordonez *et al*. 2013; Creasey *et al*. 2014).

Following the initiation of silencing via *trans*-acting small RNAs, TE silencing can be maintained by stable propagation of CG and CHG methylation, as well as reinforcement via 24 nt small RNAs derived from Pol IV transcripts that are tethered to the target gene via a Pol V transcript (Matzke and Mosher 2014; Holoch and Moazed 2015). Maintenance of silencing in the germ line is enhanced via small RNA-mediated transcriptional silencing in lineages adjacent to but distinct from the germinal lineage (Martinez and Slotkin 2012). This results in a recapitulation of the initial silencing event, in which expression of otherwise inactive elements triggers production of *trans*-acting small RNAs that are then thought to be transported to the germinal lineage (Slotkin *et al*. 2009; Li *et al*. 2010; Creasey *et al*. 2014). The net effect of this process is that active TEs can be recognized and silenced, and potentially active TEs can be kept in a stably silenced state over long periods of time.

The initiation and maintenance of TE silencing is particularly well characterized in the *Mutator* system in maize, primarily because the autonomous regulator of the system can be heritably silenced by a *trans*-acting locus called *Mu killer* (*Muk*), a naturally occurring derivative of *MuDR* that expresses a long hairpin transcript (Slotkin *et al*. 2003; Slotkin *et al*. 2005). This transposon system is composed of several related classes of cut and paste elements, all of which share similar, ∼200 bp terminal inverted repeats (TIRs) but each of which carries unique internal sequences (Lisch 2002). The system is regulated by autonomous *MuDR* elements, which carry two genes: *mudrA*, which encodes the putative transposase MURA, and *mudrB*, which encodes the helper protein MURB (Figure 1) (Hershberger *et al*. 1995; Lisch *et al*. 1999). Expression of *mudrA* and *mudrB* is convergent, with transcripts from each gene originating from within the 220 bp TIRs adjacent to each gene and extending towards the middle of the element (Figure 1). In our lines, activity of *MuDR* is monitored by a reporter element, a non-autonomous *Mu1* element located in the *a1-mum2* allele of the *A1* gene, whose function is required for color expression in both the plant and the seed (Chomet *et al*. 1991). In the seed, *Mu1* excises from *a1-mum2* somatically in the presence of a functional *MuDR* element, giving rise to spotted kernels. In the absence of functional *MuDR* elements, the kernels are uniformly pale.

**Figure 1.**
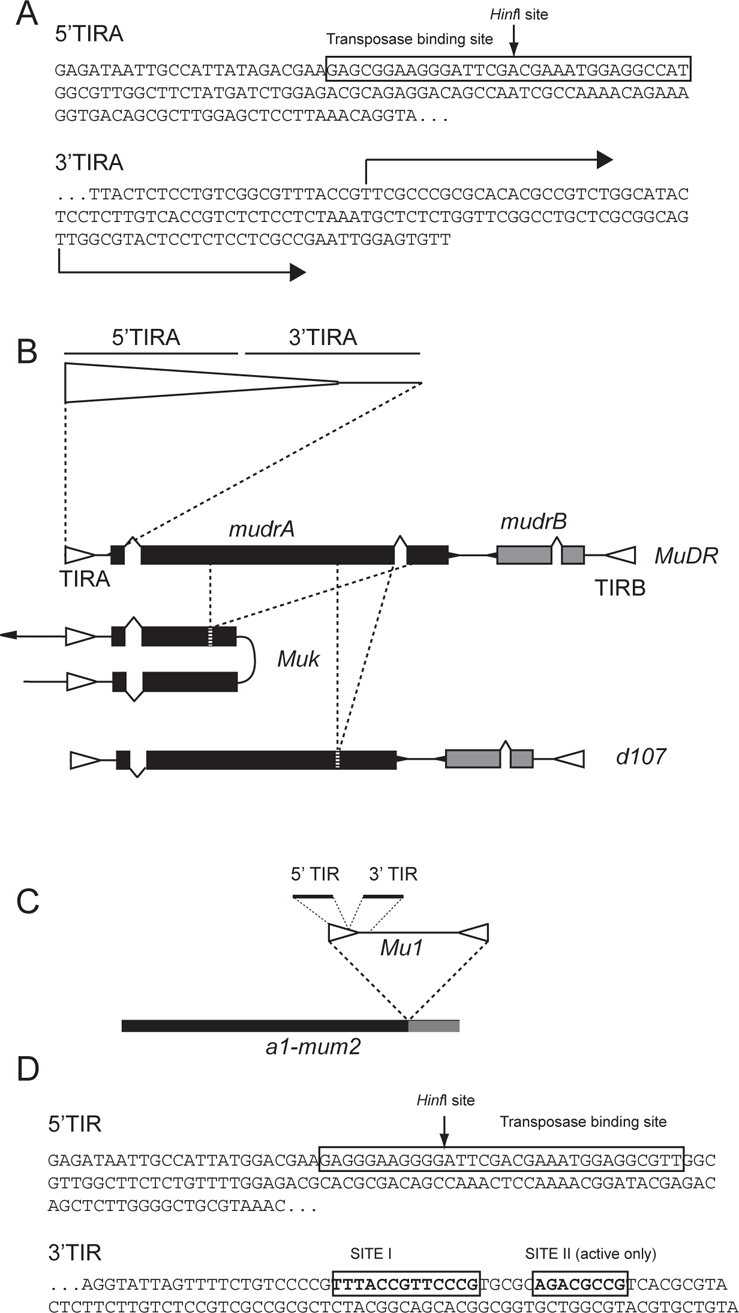
The structures of *Mu* elements examined. A) The DNA sequence of the terminal inverted repeat (TIR) adjacent to the *mudrA* gene in *MuDR* (TIRA). 5’TIRA refers to the first 144 bp of the TIR and includes the known binding site for the *MuDR* transposase. The *Hinf*I site has long been diagnostic for methylation in TIRs. The adjacent 3’TIRA includes the last 75 bp of the TIR along with 56 bp of internal sequences corresponding to a portion of the *mudrA* 5’ UTR, indicated by arrows. This region includes both of the alternative transcriptional start sites for *mudrA.* B) A diagram of the structure of *MuDR*, *Muk* and *d107*. TIRs are indicated as open triangles and exons as shaded boxes. The regions missing in *Muk* and *d107* are indicated by dashed lines. C) The structure of the *Mu1* insertion at *a1-mum2*. D) The sequence of the *Mu1* TIR, divided into its 5’ and 3’ parts. The transposase binding site is as indicated, as are additional protected sites within the 3’ portion of the *Mu1* TIR.

Also present in this and likely all maize lines are *MuDR* derivatives called *hMuDR* elements (Rudenko and Walbot 2001). Although nearly identical to portions of *MuDR*, none of these elements are intact and they do not appear to contribute to *Mutator* activity, either positively or negatively (Lisch and Jiang 2008). They are, however, a source of nuclear localized transcript and are thus the likely source of the abundant *MuDR*-similar small RNAs that have been observed in immature ears and embryos (Rudenko *et al*. 2003; Woodhouse *et al*. 2006; Nobuta *et al*. 2008). Finally, the reference maize genome contains a limited number of non-autonomous *Mu* elements with high homology within their TIRs to known active *Mu* elements (Lisch 2015). The available data suggests that both *hMuDRs* and non-autonomous elements are targeted by 24-26 nt small RNAs that are dependent on the RNA-directed DNA methylation pathway (Nobuta *et al*. 2008; Hale *et al*. 2009).

Active *MuDR* elements can be heritably and reliably silenced by genetically combining them with *Mu killer* (*Muk*), whose transcript is identical to a portion of the *mudrA* gene, as well as its associated TIR (TIRA) (Figure 1). In active *MuDR* elements, TIRA is devoid of DNA methylation. In contrast, in plants carrying both *Muk* and *MuDR*, TIRA sequences are densely methylated in all three sequence contexts, CG, CHG and CHH (where H represents any nucleotide but guanine) (Li *et al*. 2010). This methylation and the associated silencing of the *MuDR* element, can be heritably propagated over many generations, even in the absence of *Muk* (Slotkin *et al*. 2003).

Interestingly, we have found that silencing of *MuDR* by *Muk* is sensitive to changes in components of the *trans*-acting silencing (tasiRNA) pathway. Specifically, we found that a transient loss of expression of *Leafbladeless1* (*lbl1*), the maize homolog of SGS3, in leaves that emerge during the transition from juvenile to adult growth is associated with an alleviation of transcriptional silencing of the *mudrA* gene (Li *et al*. 2010). Since SGS3 works in conjunction with RDR6 to produce secondary double-stranded RNAs (Kumakura *et al*. 2009), this suggests that silencing of *MuDR* elements in leaves requires production of secondary dsRNAs triggered by small RNAs produced by the hairpin *Muk* transcript.

Here we show that cytosine methylation of different regions within the *MuDR* TIR has distinct causes and consequences, and corresponds to distinct populations of small RNAs derived from the *Muk* hairpin, the *mudrA* transcript, and other *Mu* elements in the maize genome. In addition, we demonstrate that although active *MuDR* elements can reverse methylation at one end of the TIR of a silenced *MuDR* element, they do not heritably reactivate that silenced element, nor does the silenced element inactivate the active element. Finally, we demonstrate that the previously described transient relaxation of *Muk*-induced silencing of *MuDR* during vegetative change (Li *et al*. 2010) is associated with a dramatic reduction in small RNAs targeting that element.

## RESULTS

### The absence of transposase results in default methylation of cytosines in all sequence contexts at TIRs of non-autonomous elements

When *Mutator* activity is lost due to silencing or genetic segregation of autonomous *MuDR* elements, methyl-sensitive sites within the TIRs of non-autonomous *Mu* transposons such as *Mu1* at *a1-mum2* become methylated (Chandler and Walbot 1986; Chomet *et al*. 1991). This methylation is fully reversible when a source of transposase is added, and invariably occurs when transposase is lost. Indeed this can occur in somatic sectors in developing plants when spontaneous deletions within *MuDR* elements occur, suggesting that the RNA directed DNA methylation (RdDM) pathway is competent to trigger *de novo* methylation of *Mu* elements during somatic development (Chomet *et al*. 1991; Lisch and Jiang 2008). Recent work has demonstrated that this default methylation requires a component of the RdDM pathway, *MOP1,* a protein that is homologous to *Arabidopsis* RNA-DEPENDENT RNA POLYMERASE II (Lisch *et al*. 2002; Alleman *et al*. 2006; Woodhouse *et al*. 2006). This has led to the suggestion that the TIRs of non-autonomous elements are subject to a default methylation pathway that operates in the absence of the transposase but that can be blocked and even reversed by the presence of the transposase (Hershberger *et al*. 1995; Lisch *et al*. 1995; Benito and Walbot 1997). However, this conclusion has been based on a limited number of restriction enzyme sites within the TIRs of non-autonomous elements. We wanted to understand the distribution and nature of this methylation more fully, so we examined methylation of the TIR of the non-autonomous *Mu1* element inserted into the *A1* gene in the *a1-mum2* (O’Reilly *et al*. 1985; Chomet *et al*. 1991) allele using bisulfite sequencing.

The results were entirely consistent with previous observations. In the absence of transposase, the *Mu1* TIR is extensively methylated in all three sequence contexts, although it is interesting to note that methylation of this TIR is much higher in the 5’ end of the TIR (62% methylated cytosines in the first 110 nt) than the 3’ end of the TIR (11% methylated cytosines in the second 110 nt), only marginally more than the 3% observed in the presence of the transposase (Figure 2). The 5’ end of the TIR contains sequences known to bind the MURA protein (Benito and Walbot 1997); the 3’ end of the TIR has two binding sites for unknown proteins that were previously identified (Zhao and Sundaresan 1991). Methylation was restricted to the *Mu1* TIR and did not extend into adjacent *A1* promoter sequences. In plants carrying an active *MuDR* element, nearly all of the *Mu1* methylation was lost, indicating that the presence of the transposase is sufficient to remove that methylation.

**Figure 2.**
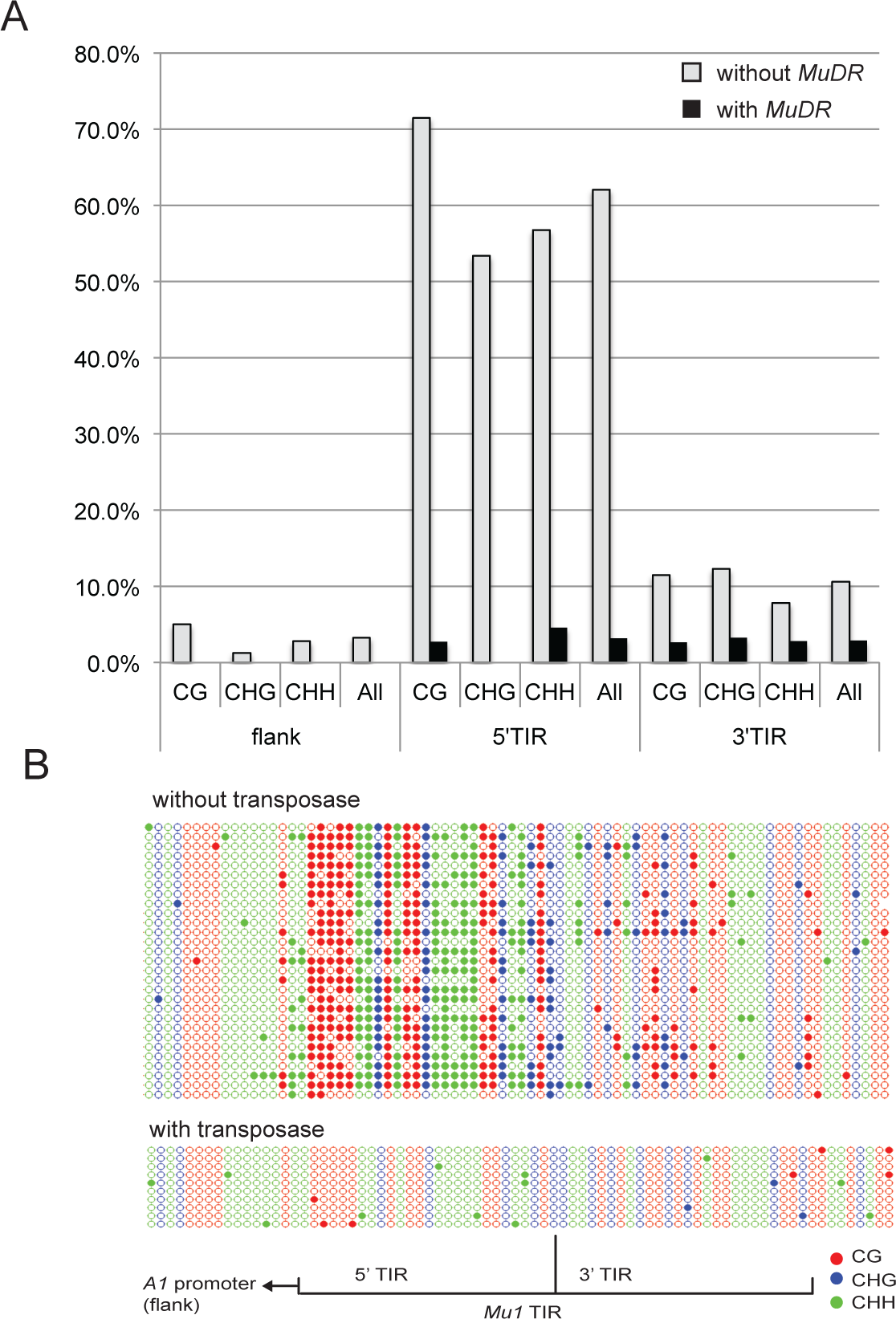
Methylation of the non-autonomous *Mu1* element in the presence and absence of an active *MuDR* element. A) Percent methylated cytosines in sequences flanking the *Mu1* insertion, as well as the 5’ and 3’ portions of the *Mu1* TIR. Cytosines are classified by sequence context, with “H” meaning any nucleotide except guanine. B) A graphic depiction of the same patterns of cytosine methylation. For this and all subsequent depictions of cytosine methylation, filled circles indicate methylated cytosines in the CG (red) CHG (blue) and CHH (green) sequence contexts. Open circles represent unmethylated cytosines. Each line represents one clone from a PCR amplification of a bisulfite-treated DNA sample.

Because the *Mu1* TIR is not identical to that of *MuDR* (roughly 83% identity over 200 bp, with the highest degree of identity −88% - within the first 100 bp), we wanted to examine the effects of the transposase on a non-autonomous element with TIRs that are identical to those of *MuDR.* Fortunately, we had available a direct derivative of *MuDR* (*MuDR-d107, or d107*) that is identical to the autonomous element with the exception of a 700 bp deletion within a conserved portion of the *mudrA* transposase gene (Lisch 1995;

Lisch and Jiang 2008) (Figure 1). As is the case for other *MuDR* deletion derivatives, sequence analysis of the deletion in *d107* suggests that it arose as a consequence of strand slippage during gap repair following excision of *MuDR(p1)* (Figure S1) (Hsia and Schnable 1996). This derivative cannot cause excision of the reporter element, nor can it trigger hypomethylation of non-autonomous elements, suggesting that it does not make a functional transposase. However, it is transcriptionally active, producing a full-length *mudrB* transcript and a polyadenylated but internally deleted *mudrA* transcript. *d107* also has the advantage of being at the same chromosomal location as the originally cloned *MuDR* element at position 1 on chromosome 3L (*p1*) (Chomet *et al*. 1991) and can be efficiently silenced by *Muk* (Slotkin 2005). Thus, the only difference between *d107* and the functional *MuDR* from which it was derived is the presence of the deletion.

Examination of the TIR of *d107* revealed that the default methylation that we observed at *Mu1* also occurs within the *d107* TIR. As in the case of *Mu1*, methylation was largely restricted to the 5’ end of the TIR (Figure 3B). With this in mind, we have split analysis of this TIR into 5’ (5’TIRA) and 3’ (3’TIRA) portions (Figure 1A). 5’TIRA includes the first 144 bp of TIRA. This region of the TIR includes the binding site for the transposase (MURA) (Benito and Walbot 1997) and is the most highly conserved region among *Mu* elements (Bennetzen 1996). 3’TIRA includes the last 75 bp of the TIR along with 69 bp of internal sequences corresponding to a portion of the *mudrA* 5’ UTR. This region includes both of the alternative transcriptional start sites for *mudrA* (Hershberger *et al*. 1995).

**Figure 3.**
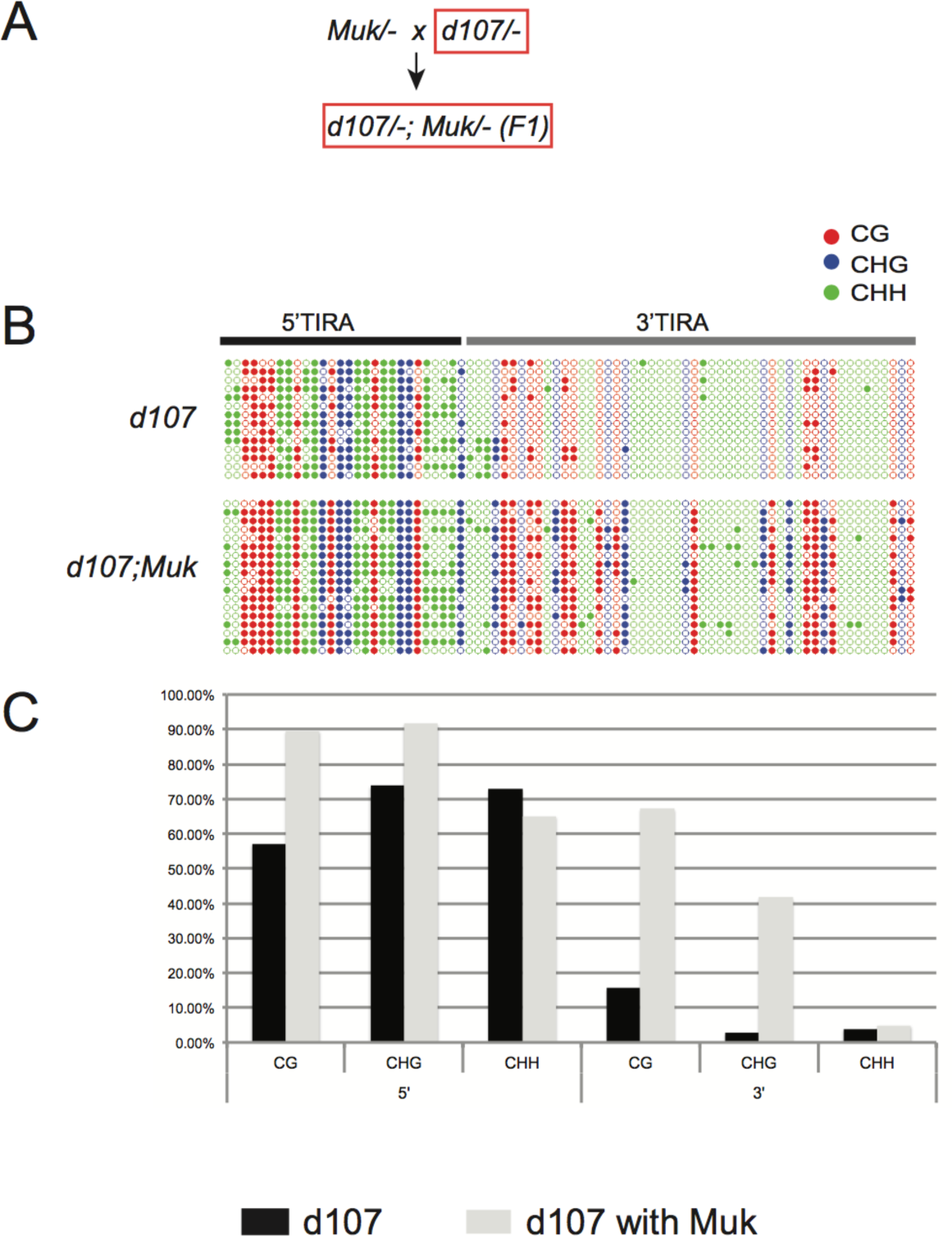
Depiction of patterns of methylation in TIRA of the deletion derivative, *d107*. A) The cross used to generate the individuals examined (one of each genotype) (boxed red) B) A graphic representation of the patterns of methylation at *d107* TIRA. Samples include a transcriptionally active *d107* and *d107* in the presence of *Muk.* Cytosines in different sequence contexts (CG, CHG and CHH) are as indicated. C) Percent methylation of cytosines in each sequence context in the same samples depicted graphically, with the results separated into 5’TIRA and 3’TIRA.

Within transcriptionally active *d107*, 68% of the cytosines in 5’TIRA were methylated. In contrast, only 7% of the cytosines in 3’TIRA were methylated (Figure 3B; *P* < 0.0001, Fisher’s exact test). These data indicate that the default methylation observed at *Mu1* is also observed at *d107*. However, because *d107* is transcriptionally active, we can conclude that the dense methylation we observe in 5’TIRA in all sequence contexts in this derivative is not sufficient to trigger transcriptional silencing.

In order to determine the effects of *Muk* on *d107* a plant carrying *d107* was crossed to a plant carrying *Muk* and the pattern of methylation at TIRA was examined in a progeny plant carrying both *d107* and *Muk* (Figure 3A). In this plant, the level of methylation of 5’TIRA was quite similar to that observed in *d107* not exposed to *Muk* (78% and 69%, respectively) (*P* = 0.2327, Fisher’s exact test). In contrast, exposure to *Muk* caused extensive CG and CHG methylation of 3’TIRA of *d107* (30%), significantly higher than was observed in *d107* by itself (7%) (*P* < 0.0001, Fisher’s exact test). This pattern of methylation is nearly identical to that seen in TIRA of *MuDR* elements as they are being silenced by *Muk* in immature ears (Li *et al*. 2010).

Interestingly, we find that the sequence context of the methylated cytosines is quite different between 5’TIRA and 3’TIRA of *d107*. In both *d107* and in F1 *d107*;*Muk* plants the cytosines are densely methylated in all three sequence contexts in 5’TIRA. In contrast, in 3’TIRA, cytosine methylation in the CHH context was uniformly low (4% for *d107* and 5% for *d107*;*Muk*) ( *P* = 0.6210, Fisher’s exact test), but CG and CHG was considerably enriched in *d107;Muk* relative to *d107* plants in 3’TIRA (67% versus 16% for CG (*P* < 0.0001, Fisher’s exact test), and 42% versus 3% for CHG, respectively (*P* < 0.0001, Fisher’s exact test). Collectively, from the examination of *d107*, we conclude that methylation of 5’TIRA is independent of transcriptional silencing, and that exposure of transcriptionally active *d107* to *Muk* induces new CG and CHG methylation to the 3’TIRA, the region that includes both alternative transcriptional start sites. These data suggest that 5’TIRA and 3’TIRA methylation may involve distinct molecular mechanisms that result in distinct patterns of cytosine methylation.

### Active elements remove default methylation within silenced elements but do not heritably reactivate them

Having established the nature of default methylation, and the distinction between default methylation and methylation induced by *Muk* at *d107*, we sought to examine the effects of an active element on a previously silenced element. To do this, we crossed a plant carrying *MuDR* at position 1 (henceforth referred to as “p1”) to a plant carrying *Muk*. Previous analysis had determined *MuDR* TIRA is densely methylated in the mature leaves and immature ears of F1 (p1/-;*Muk/-*) plants (Li *et al*. 2010). Plants carrying p1 and *Muk* were crossed to plants carrying an active *MuDR* element at a second chromosomal position (referred to as “p5”) (Figure 4A) (Singh *et al*. 2008). A progeny plant carrying p1 that had been silenced in the previous generation but that no longer carried *Muk* (referred to as p1*) was then compared to a sibling carrying both p1* and p5 in order to examine the heritability of silencing and the effects of an active element on a silenced element. Consistent with the fact that *Muk* induces heritable silencing of *MuDR* elements, p1* by itself was extensively methylated. Overall, 5’TIRA of p1* had 81% methylated cytosines and 3’TIRA had 48%, similar to the levels of methylation in F1 (p1/-;*Muk/-*) plants (Li *et al*. 2010). These data confirm our previous observation that methylation at 5’TIRA established due to the presence of *Muk* is maintained in subsequent generations in its absence (Li *et al*. 2010). Analysis of a sibling plant that carried both p1* and p5 revealed extensive changes in the pattern of methylation at p1* TIRA. In the 5’TIRA of these plants, only 15% of cytosines were methylated, suggesting that the methylation established in this region due to the activity of *Muk* was largely lost in a manner similar to what is observed at non-autonomous elements when exposed to the transposase. In contrast, within 3’TIRA 34% of the cytosines remained methylated in this region, somewhat less than the 48% methylation observed in 3’TIRA in siblings that carried only p1* and F1 *MuDR;Muk* parent in the previous generation (44%) (*P* = 0.0032, Fisher’s exact test).

**Figure 4.**
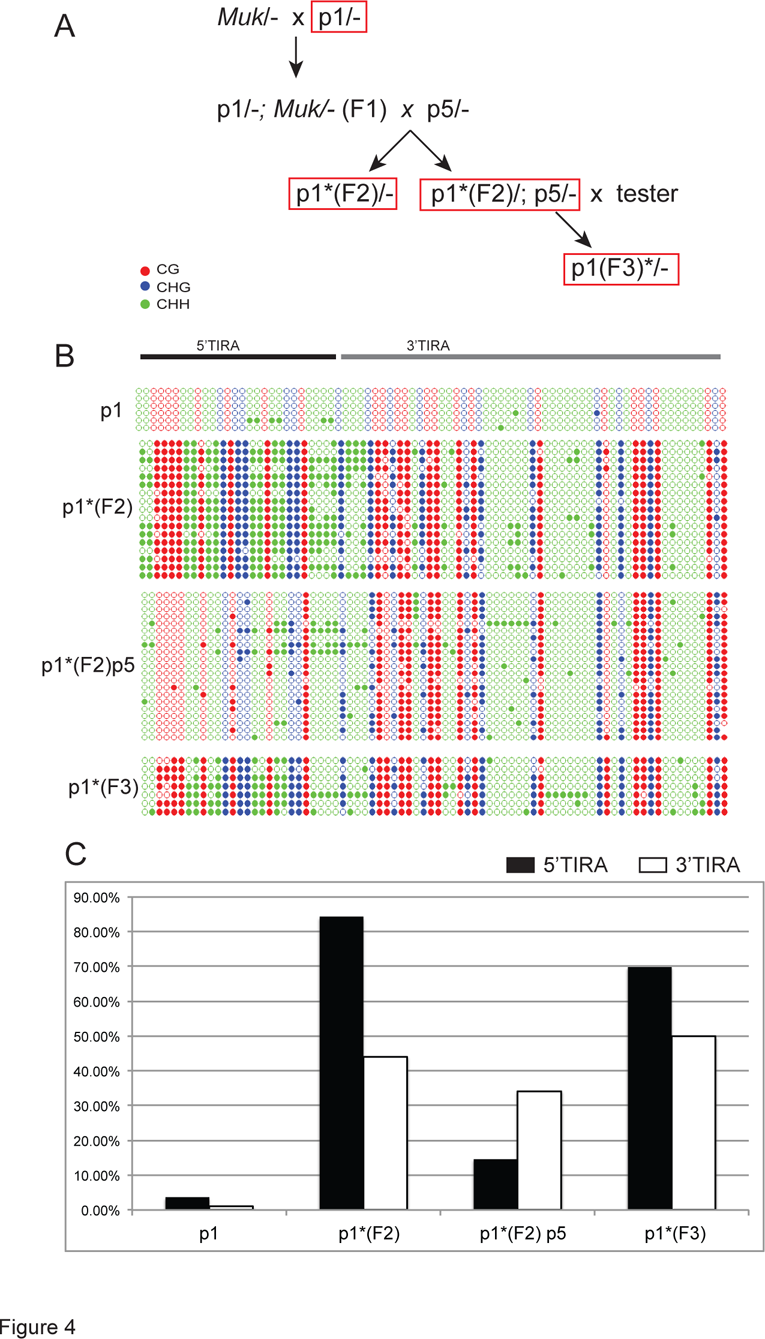
The effect of an active element on methylation of a silenced element. A) The crosses used to generate the samples subject to bisulfite sequencing (boxed red) B) A graphic representation of patterns of cytosine methylation within TIRA in plants of the indicated genotypes. Cytosines in different sequence contexts (CG, CHG and CHH) are as indicated. C) Percent methylation of cytosines within 5’ TIRA and 3’TIRA in plants of the indicated genotype.

Having established that an active element can reverse at least some of the methylation associated with *Muk*-induced silencing of p1, we wanted to determine whether or not this has a heritable effect on the silenced element. In order to do this, a plant carrying p1* by itself and a sibling carrying both p1* and p5 were test crossed to a plant that lacked both *MuDR* and *Muk,* referred to as an *a1-mum2* tester (Figure 4A).

When crossed to the *a1-mum2* tester, the plant carrying only p1* gave rise to an ear all of whose kernels were uniformly pale. The plant carrying both p5 and p1* yielded an ear that segregated for a single active element (44 spotted to 41 pale kernels), suggesting that p5 had not been silenced by p1*, and p1* had not been heritably activated as a consequence of exposure to p5. Analysis of methylation of TIRA in a progeny plant that carried only p1* but that had been exposed to p5 in the previous generation (p1*/-F2 in Figure 4A) revealed that exposure to p5 had no obvious heritable effect on the silenced element. 70% of the cytosines in 5’TIRA were methylated, as were 50% of the cytosines in 3’TIRA (Figure 4C). Further, when this plant was test crossed, none (0/137) of its progeny kernels were spotted.

In order to extend this observation, we performed a cross between a plant carrying a previously silenced p1 element (p1*) by a plant that carried an active *MuDR* element at a third position, p4 (Singh *et al*. 2008). The resulting ear segregated for a single active *MuDR* element (52% spotted progeny kernels). Spotted and pale progeny kernels were genotyped for p1 and p4 and then test crossed to plants that lacked either element. Of the plants grown from spotted progeny kernels, six of fourteen carried p1* and all of them contained p4. When test crossed, these fourteen plants gave rise to an average of 52% spotted progeny (Supplemental Table 1), consistent with the segregation of a single active element. Siblings grown from spotted kernels that lacked p1* gave rise to an average of 50% spotted progeny. Of the pale progeny, eight of seventeen carried p1*.

When test crossed, none of these gave rise to any spotted progeny. From these experiments, we conclude that the presence of the transposase from the active element had no heritable effect on the silenced element, nor did the silenced elements affect the active element.

### 22 nt *mudrA*-specific small RNAs are associated with silencing of *MuDR* by *Muk*

Previously, we had demonstrated that small RNAs are associated with silencing of *MuDR* by *Muk* (Slotkin *et al*. 2005). This conclusion was based on gel hybridization of RNAs from young juvenile leaves. In order to more comprehensively characterize these small RNAs, small RNAs were sequenced from plants lacking both *Muk* and *MuDR*, plants carrying a single active *MuDR* element, plants carrying only *Muk*, and plants carrying both *MuDR* and *Muk* (F1 plants). In each of these cases, tissue was collected from young leaf 2. Leaf 2 was chosen because it is the tissue that had previously shown ample evidence of an accumulation of *Muk*-specific small RNAs, and because TIRA is heavily methylated in all three sequence contexts in F1 plants in this leaf (Li *et al*. 2010).

Further, in contrast to immature ears, this leaf also lacks the ubiquitous *MuDR*-homologous heterochromatic small interfering RNAs (hc-siRNAs) present in all genotypes regardless of activity present in that tissue (Woodhouse *et al*. 2006). Small RNAs of leaf 6 from F1 plants were also analyzed because previous work in our laboratory had demonstrated that this leaf exhibits a striking loss of TIRA methylation, concomitant with a loss of expression of *LBL1,* an important component of the tasiRNA silencing pathway in maize (Li *et al*. 2010; Dotto *et al*. 2014).

Consistent with previous results, plants with *Muk* by itself contained large numbers of *MuDR*-identical small RNAs, the vast majority of which were 21 and (particularly) 22 nucleotides in length (Figure 5 and Supplementary Table 2). These small RNAs were oriented in both sense and antisense orientation relative to *mudrA*, and were restricted to the portion of the *Muk* transcript that can form an inverted repeat, consistent with the hypothesis that the small RNAs are a product of processing and/or amplification of the hairpin transcript produced by *Muk*. In contrast, plants carrying only *MuDR* and those that lacked both *MuDR* and *Muk* had very few *MuDR*-identical small RNAs.

**Figure 5.**
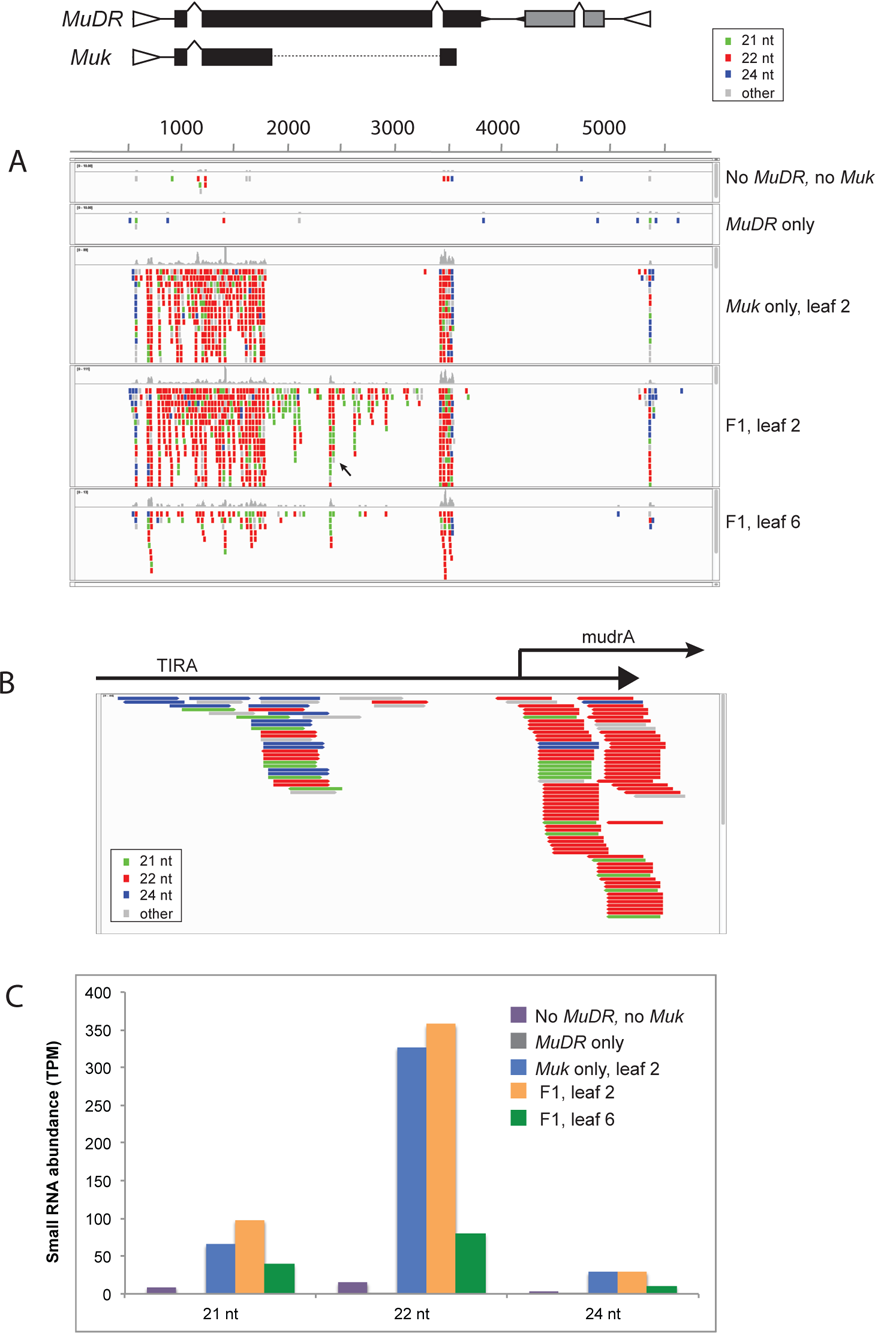
Small RNAs associated with silencing of *MuDR* by *Muk*. A) A representation of perfectly matched small RNAs from tissues of various genotypes mapped onto *MuDR*. A map of the regions present in *Muk* is provided for reference. Small RNAs are color coded for size, as indicated. Note that small RNA samples from plants containing only *MuDR* or neither *MuDR* nor *Muk* have very few small RNAs matching *MuDR*. B) Small RNAs with exact matches to the first 250 bp of *MuDR* (data is taken from F1 p1;*Muk* leaf 2). C) Numbers of perfect matches to *MuDR* of the indicated size classes to the indicated genotypes.

The presence of a deletion within *Muk* relative to *MuDR* resulted in a junction that is unique to *Muk*. We found one prominent group of 22 nt small RNAs that spanned this junction and thus can only be derived from *Muk* (Figure S2). Interestingly, this particular class of unique, *Muk*-specific small RNAs are largely in the sense orientation relative to the *mudrA* transcript, suggesting that processing of the *Muk* hairpin transcript at this site favors retention of small RNAs in this orientation.

The small RNAs sequenced from plants carrying both *MuDR* and *Muk* are quite similar to those observed in plants carrying only *Muk*, with one important exception. In these plants there are many 21 and 22 nt small RNAs corresponding to regions of *MuDR* that are not in present in *Muk* (Figure 5A). Although these small RNAs are in both orientations relative to *mudrA*, a prominent cluster, mapping approximately 1880 nt from the 5’ end of *MuDR* is primarily in the antisense orientation (Figure 5A, arrow).

Given that these *mudrA*-specific small RNAs are only present when both *MuDR* and *Muk* are combined, these data suggest a transitive process by which *mudrA* transcript is converted into double-stranded RNA and then cleaved into 21 and 22 nt small RNAs, presumably triggered by the presence of small RNAs generated by *Muk*. Interestingly, the ratio of 22 nt to 21 nt small RNAs differs between the regions shared by both *MuDR* and *Muk* and the regions only present in *MuDR*. In the former, the ratio of 22 nt to 21 nt small RNAs is 5.1:1. In the latter it is 1.4:1 (*P* < 0.0001, χ^2^ test) (Supplemental Table 3).

Given that these distinct size classes require distinct DCL proteins, generally DCL4 for 21 nt *trans*-acting small RNAs and DCL2 for the 22 nt size class (Schwab and Voinnet 2010), this observation suggests the relative contribution of these two proteins may differ in these two distinct regions.

Interestingly, the vast majority of small *MuDR*-identical small RNAs in plants that carry only *Muk* or that carry both *MuDR* and *Muk* correspond to transcribed *mudrA* sequences. This is most apparent at the transcriptional start site of *mudrA*, where there is a prominent cluster of 22 nt small RNAs, most of which are in an antisense orientation relative to *mudrA* (Figure 5B). There are also a smaller number of small RNAs corresponding to sequences upstream of the start of transcription within TIRA. However, many of them are sizes other than 22 nt. Further, when two mismatches are permitted, roughly half of them have sequences that match a non-autonomous *Mu* element present elsewhere in the maize reference genome rather than *MuDR* (Figure S3) (Tan *et al*. 2011).

The distribution of these two distinct small RNA populations in all of the plants carrying *Muk* matches the distribution of DNA methylation within TIRA of both *d107* and silenced *MuDR* elements. The variably sized and polymorphic small RNAs correspond to the 5’ end of TIRA that is default methylated in all sequence contexts in the absence of the transposase in *d107* (Figure 3). The more numerous 22 nt small RNAs correspond to the expressed portion of TIRA whose stable CG and CHG methylation is triggered in response to *Muk* (Figures 3 and 4). Interestingly, analysis of large populations of small RNAs from *mop1* mutant and wild type immature ears, as well as *lbl1* mutant and wild type leaf apexes provides similar evidence (Supplemental Table 4). In these libraries, derived from plants that did not carry *MuDR* or *Muk,* there are very few small RNAs of any size class perfectly matching either 5’TIRA or 3’TIRA. When two mismatches are permitted, there are a substantial number of small RNAs matching TIRA, but the vast majority of them are in 5’TIRA. In combination with our analysis, these data suggest that *de novo* methylation of 5’TIRA, but not 3’TIRA, is mediated by “background” small RNAs with one or more mismatch, but de novo methylation of 3’TIRA requires 21 and 22 nt small RNAs derived from the *Muk* hairpin and processed Pol II *mudrA* transcript.

Given the involvement of *LBL1* in the production of dsRNA, and the requirement for *LBL1* in methylation of TIRA in leaf tissue, we hypothesized that there would be a reduction of small RNAs in leaves that are transitional between juvenile and adult, which exhibit a loss of both TIRA methylation and a reduction of *Lbl1* expression (Li *et al*. 2010). In fact we observed a dramatic reduction in the number of small RNAs of all size classes in transition leaves (Figure 5C and Supplemental Table 2).

### TIRB silencing is not associated with small RNAs

*Muk* does not include sequences from *mudrB*. However, it does include TIRA, which is 99% identical over the first 180 bp with TIRB, suggesting that small RNAs from the *Muk* hairpin transcript would be hypothetically competent to direct methylation of TIRB. Although *mudrB* is eventually silenced by *Muk,* the trajectory of *mudrB* silencing is distinct. Unlike *mudrA*, which is transcriptionally silenced by the immature ear stage in *p1*;*Muk* F1 plants, *mudrB* is not transcriptionally silenced by this stage (Slotkin *et al*. 2003). Instead, in these plants transcript from *mudrB* is readily detectable, but it does not appear to be polyadenylated. By the next generation, however, plants that carry only *p1*\* do not have detectable *mudrB* transcript. Further, *Muk* can only heritably silence *mudrB* in this way when this gene is in *cis* to *mudrA* (on the same transposon); it is not silenced when it is in *trans* to a *mudrA* gene that is being silenced (Slotkin *et al*. 2005).

Previous work using RNA gel hybridization showed that small RNAs similar to *mudrB* did not accumulate to high levels in F1 *MuDR*;*Muk* plants (Slotkin *et al*. 2005). In some ways, this was surprising given that TIRB is identical over much of its length to TIRA, and the hairpin formed by the *Muk* transcript includes all of TIRA (and thus TIRB as well). A closer examination of the small RNA population in F1 leaf 2 tissues provides a possible explanation for this discrepancy. Although the two TIRs are quite similar to each other, they are more diverged near the internal portion of each TIR, within 3’TIRA and 3’TIRB (Figure 6C). The *mudrA* transcript initiates 168 bp from the end of the element and the *mudrB* transcript initiates 163 bp from the other end of the element (Hershberger *et al*. 1995). Since this is the region in which TIRA and TIRB begin to diverge in sequence, there are very few 22 nt small RNAs that perfectly match TIRB, particularly in the transcribed portion of TIRB (Figure 6A and B). If silencing requires the presence of both the target transcript and small RNAs from the *Muk* hairpin, this distribution of small RNAs may explain why *Muk* acts only indirectly on *mudrB* via a distinct mechanism.

**Figure 6.**
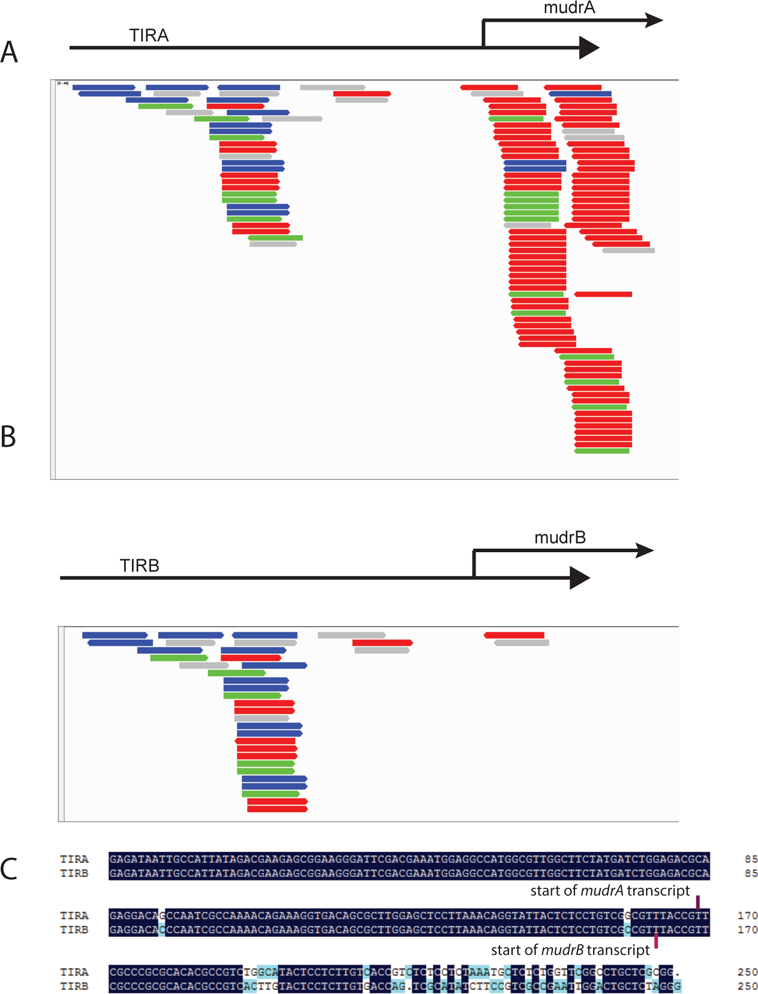
Small RNAs in F2 leaf 2 plants with perfect matches to TIRA (A) and TIRB (B). C) An alignment of TIRA and TIRB with the transcriptional start sites for *mudrA* and *mudrB* as indicated.

## Discussion

The evidence presented here suggests that two distinct silencing pathways operate on *MuDR* TIRA as it is being silenced. One pathway, dependent on MOP1, appears to involve a number of heterogeneous small RNAs likely derived from other *Mu* TIRs in the genome that target 5’TIRA for DNA methylation in the absence of the transposase. They do not, however, trigger transcriptional silencing, as is evidenced by *d107*, which accumulates dense methylation in 5’TIRA but is transcriptionally active. In contrast, when *d107* or *MuDR* is silenced by *Muk*, methylation spreads from 5’TIRA into the transcribed portion of the *mudrA* (Figure 3 and (Li *et al*. 2010)). Once *MuDR(p1)* is silenced, a source of transposase can reverse the methylation at 5’TIRA but does not reverse the heritably silenced state of *MuDR(p1)*, nor does it reverse methylation within 3’TIRA (Figure 4). Thus, one can gain methylation at 5’TIRA and not gain transcriptional silencing, and one can lose 5’TIRA methylation and not lose heritable transcriptional silencing. Interestingly, these data also suggest that the methylation required for a heritably silenced state is restricted to the 3’TIRA, and is largely composed of CG and CHG methylation (Figure 4).

The second pathway involves 21 and 22 nt small RNAs specifically associated with sequences within 3’TIRA corresponding to the transcribed portion of *mudrA*. These small RNAs correspond to cytosines within 3’TIRA that are stably methylated in the CG and CHG sequence contexts, and are only observed in leaves carrying *Muk*, either by itself or in combination with *MuDR*. Plants carrying both *MuDR* and *Muk* also have many small RNAs that are derived from portions of *mudrA* that are not present in *Muk*. This is consistent with the activity of a transitive process that is triggered by small RNAs processed from the *Muk* hairpin. This hypothesis is supported by the observation that in leaf 6 in which *lbl1* expression is reduced, the number of these small RNAs is also reduced (Figure 5, Table S1) (Li *et al*. 2010). The fact that the 22 nt small RNAs do not extend upstream of the transcriptional start site of *mudrA* suggests that these small RNAs are processed almost entirely from transcript initiated from within TIRA rather than the double-stranded hairpin formed by *Muk*, which includes 5’TIRA. Since *LBL1* works in conjunction with *RDR6* to produce a variety of *trans*-acting small RNAs, these data suggest that the *Muk* triggers silencing in leaves via a pathway that interacts with Pol II TE transcripts in a manner that is similar to that described in *Arabidopis* (Wu *et al*. 2012; Nuthikattu *et al*. 2013; Panda and Slotkin 2013; Panda *et al*. 2016). In this context, an interaction between small RNAs and Pol II transcript represents a natural and expected means by which otherwise active TEs are recognized and silenced. The key here is the source of small RNAs that can trigger silencing. For high copy number elements, a low level of expression of aberrant transcripts in germinal lineages could be sufficient to trigger silencing of transiently active elements. For low copy number elements such as *Mu* or *Ac*, it may be necessary for rearrangements to arise prior to effective silencing. Interestingly, however, maize lines containing larger numbers of *Mu* elements are prone to spontaneous silencing (Robertson 1986; Skibbe *et al*. 2012). This phenomena appears to be a function of copy number, as single copy number *MuDR* line, which was derived from such a line does not show evidence of spontaneous silencing, and if copy number is allowed to increase in this line spontaneous silencing begins to occur (D. Lisch, unpublished). We speculate that this spontaneous silencing could be due to the accumulation of aberrant *MuDR* elements that collectively trigger silencing of active elements in this line.

One important conclusion from our analysis is that default methylation such as that observed at *d107* and *Mu1* is not sufficient to trigger silencing. *d107* expresses a transcript and *Mu1* element has a functional outward reading promoter even when it is methylated (Barkan and Martienssen 1991). Because hypomethylated *Mu1* elements can be re-methylated due to genetic segregation of *MuDR*, or even in somatic sectors in which the transposase is lost, it is apparent that this pathway is competent to trigger *de novo* methylation during plant development (Lisch *et al*. 1995; Lisch and Jiang 2008). However, this methylation is readily reversed when transposase is reintroduced. This may be because binding of the transposase to the TIR directly blocks methyltransferase activity or due to demethylation activity on the part of the transposase as has been proposed for the *Spm* transposase (Cui and Fedoroff 2002). These results are reminiscent of patterns of methylation in other transposable elements. For instance, although active maize *Spm* elements are extensively methylated throughout much of the element, they are hypomethylated in the region immediately upstream of their transcriptional start site. In contrast, silenced *Spm* elements are extensively methylated in this region (Banks *et al*. 1988). Further, when silenced *Spm* elements are exposed to active *Spm* elements, an adjacent region in the active element can be demethylated (Cui and Fedoroff 2002). Methylation within the *Tam3* transposon in *Antirrhinum majus* is reversible and the loss of methylation requires the presence of the transposase, suggesting that methylation of this TE may be a consequence of the loss of transposase rather than a cause of silencing (Hashida *et al*. 2006). Our data is consistent with that hypothesis, at least with respect to methylation within TIRA.

The default methylation at 5’TIRA is dependent on the maize homolog of RDR2, MOP1, which is required for the vast majority of 24 nt heterochromatic siRNAs (hc-siRNAs) (Lisch *et al*. 2002; Alleman *et al*. 2006; Nobuta *et al*. 2008). Previous work in our laboratory has demonstrated that heritable silencing of *MuDR* by *Muk* occurs efficiently in a *mop1* mutant background in the absence of those hc-siRNAs, and that *Muk*-specific small RNAs are retained in this mutant (Woodhouse *et al*. 2006). Further, in immature ears, the presence of the hc-siRNAs, likely derived from *hMuDR* elements in this genetic background, has no effect on an otherwise active *MuDR* element. Thus, it would appear that *de novo* silencing of *MuDR* elements requires small RNAs derived from Pol II transcript, but likely not those derived from processed Pol IV transcripts.

This would explain why silenced *MuDR* elements do not silence active elements, as the silenced elements would be expected to produce only Pol IV transcript, which would not be expected to produce *trans*-acting small RNAs. More generally, it suggests that the presence of previously silenced elements, by itself, is not sufficient to initiate silencing. Indeed, this has been demonstrated for reactivated elements in *Arabidopsis* (Kato *et al*. 2004). This in turn suggests that transient loss of silencing of previously silenced *MuDR* elements, such as has been observed in pollen (Slotkin *et al*. 2009) would only be expected to function to reinforce silencing if those previously silenced elements produced aberrant Pol II transcripts with a propensity to be processed into *trans*-acting small RNAs. This seems to be the case for easiRNAs in *Arabidopsis*, which are produced from aberrant hairpin micro-RNA-like transcripts that target transposons (Creasey *et al*. 2014). Thus, we suggest that the key to silencing reinforcement in tissues such as pollen is not reactivation of silenced elements *per se*, but reactivation of “zombie” elements, whose Pol II transcripts, when expressed, are competent to reinforce or trigger silencing in *trans*. These elements could be similar in structure of *Muk* and result in the production of a hairpin, but any feature of their transcript that is recognized as aberrant that would be sufficient. In our low copy *Mutator* line, it would appear that these zombie elements are not present. Thus, pollen that carries both a silenced *MuDR* element and an active element does not give rise to progeny in which the active element has been silenced, despite evidence that silenced *MuDR* elements, like other silenced transposons, are transcriptionally activated in the pollen vegetative nucleus as part of a strategy of silencing reinforcement (Slotkin *et al*. 2009; Martinez *et al*. 2016).

Our observations are consistent with a relatively simple model (Figure 7). TIRA methylation is absent whenever a functional transposase is present, presumably because the transposase blocks methyltransferase activity. When 22 nt small RNAs are introduced from the *Muk* hairpin, they trigger RDR6-dependent production of double-stranded RNA using the Pol II-derived *mudrA* transcript as a template. This double-stranded RNA is cleaved by a DICER (presumably DCL2 and/or DCL4) and the resulting small RNAs are then used to direct DNA methylation, largely in the CG and CHG sequence contexts in 3’TIRA. Meanwhile, because the transposase is no longer present, hc-siRNAs derived from other silenced *Mu* elements present in the genome can direct *de novo* methylation of 5’TIRA, exactly as they do at *d107* and *Mu1* in the absence of functional transposase (Figures 2 and 3). According to this model, processed *mudrA* transcript and the resulting 3’TIRA methylation is the cause of transcriptional silencing, and 5’TIRA methylation is a consequence of that silencing. Based on this model, we would predict that a *MuDR* derivative that had a genetic lesion that prevented it from expressing Pol II transcript would accumulate methylation in 5’TIRA but would not accumulate methylation in 3’TIRA when combined with *Muk*.

**Figure 7.**
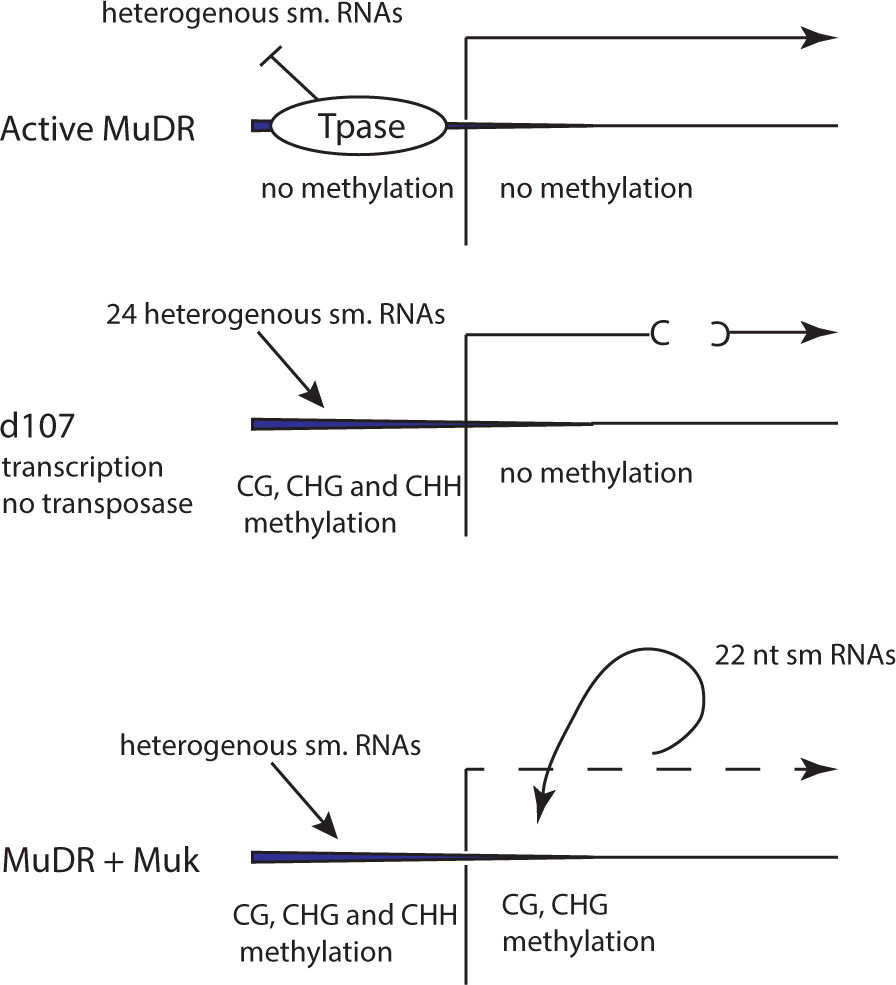
A model for the interaction between different classes of small RNAs and DNA methylation in 5’TIRA and 3’TIRA.

The involvement of two parallel pathways interacting with *MuDR* TIRA raises some interesting questions. The default methylation pathway suggests that there are small RNAs that are competent to trigger *de novo* methylation that are present even when the elements are active. However, active *MuDR* elements retain that activity in the face of this “soup” of hc-siRNAs, and the same is true for some other active transposons, such as CACTA elements in *Arabidopsis* and *Mu* elements in maize (Kato *et al*. 2004; Woodhouse *et al*. 2006). In contrast, others, such as ONSEN and *Tos17*, are rapidly re-silenced after periods of activity, a process that requires Pol IV (Hirochika *et al*. 1996; Ito *et al*. 2011; Matsunaga *et al*. 2015). Since Pol IV transcript is thought to be derived from previously methylated elements, this would suggest that at some threshold, silenced elements can contribute to *de novo* silencing of active elements. In contrast, *Muk* silencing of *MuDR* does not appear to involve 24nt Pol IV-dependent small RNAs, presumably because processing of the *Muk* hairpin can produce siRNAs in the absence of hc-siRNAs (Woodhouse *et al*. 2006). Further, those hc-siRNAs that are present in the background largely target a portion of active *MuDR* elements (5’TIRA) that is irrelevant to transcriptional gene silencing (Figure S3). This may be due to random sequence divergence within the relevant portions of TIRA relative to other *Mu* elements, or due to selection in favor of differences between autonomous and non-autonomous elements at the *mudrA* and *mudrB* transcriptional start sites. Indeed, we find few hc-siRNAs specifically targeting the transcribed portion of TIRA. Further, a more comprehensive analysis of all available small RNAs derived from a number of tissues and genetic backgrounds reveals that the vast majority of small RNAs with zero, one or two mismatches to *MuDR* map to 5’TIR rather than 3’TIR (Supplemental Table 4). The absence of small RNAs directly targeting the transcribed portions of *mudrA* and *mudrB*, despite the presence of *hMuDRs* suggests that selection may have favored sequence divergence between currently active and previously silenced autonomous elements (*hMuDRs*).

Collectively, our data suggests that active elements in maize remain active because of an absence of small RNAs that are competent to trigger heritable silencing, and that small RNAs competent to trigger silencing are most likely those that directly target the Pol II transcript arising from those active elements, rather than the small RNAs derived from previously silenced elements. Further, our data suggest that heritable silencing information is contained within 3’TIRA in the form of CG and CHG methylation. Finally, our analysis of small RNAs in transition leaves reveals that the loss of *LBL1* in these leaves results in a reduction of small RNAs targeting *mudrA*, suggesting that the accumulation of these small RNAs are dependent on the tasiRNA pathway.

## Author contributions

D.B performed the bisulfite analysis and contributed to experimental design. H.L performed the small RNA sequencing analysis contributed to experimental design. S.K. performed bisulfite analysis on the *MuDR* active individual. M.Z. performed analysis of the small RNA data set. D.L wrote the manuscript, performed the genetic analysis and analyzed the small RNA and deletion junction analysis.

## Acknowledgements

This work was supported by the National Science Foundation (Grants DBI-1237931and DBI-0820828 to D.L.) as well as the Purdue Center for Plant Biology. We thank Xinyan Zhang and Thomas Peterson for providing useful comments.

## Materials and Methods

### Plant materials

The minimal *Mutator* line consists of one fully active *MuDR* element at position 1 (p1) on chromosome 3L (Chomet *et al*. 1991). *Muk* is a derivative version of *MuDR* as described previously (Slotkin *et al*. 2005). Activity is monitored in seeds via excisions of a non-autonomous *Mu1* element inserted into the *a1-mum2* allele of the *A1* color gene (Chomet *et al*. 1991). All plants used in these experiments are homozygous for *a1-mum2.* “Test crosses” refer to crosses to plants that are homozygous for *a1-mum2* but that lack *MuDR* or *Muk*.

The crosses used to generate the genotypes examined are depicted in Figures 3A and 4A. In order to construct lines containing silenced *MuDR(p1)* elements (p1*) with and without active elements, plants carrying p1 were crossed to plants heterozygous for *Muk*. Progeny plants that were heterozygous for *Muk* and that carried p1 were then crossed to plants that carried *MuDR(p5)* (p5). Progeny that lacked *Muk* and that carried silenced p1 (p1*) with or without p5 were then compared. Plants carrying p1* and p5 were then test crossed to *a1-mum2* testers, and progeny plants carrying only p1* were examined. Tests of the ability of a second active element, p4, to activate a silenced element were performed by crossing a plant carrying p4 to a plant carrying a previously silenced p1* element and then test crossing plants carrying both p1* and p4. Progeny of this cross were then separated into spotted (exhibiting somatic excisions of the reporter element) and pale kernels, genotyped for p1 and p4 and then test crossed. Genotyping for p1 employed primers RLTIR2 and Ex1. Genotyping for p4 employed primers RLTIR2 and p4flankB. Genotyping for p5 employed primers RLTIR2 and p5flankB (all primers used in experiments described in this manuscript are supplied in Supplemental Table 5).

### Tissue sampling

All plants used in bisulfite sequencing experiments were genotyped individually. Immature ears, approximately 10 cm long, were collected from each individual plant. To check for variation in patterns of methylation, fully mature leaves (the third leaf from the top of each plant) were also examined (Figure S5). Given that the results were nearly identical for ears and leaves, only the data from ears is presented here. For small RNA analysis, the distal 10 cm of emerging leaves was collected as previously described (Li *et al*. 2010).

### Genomic Bisulfite sequencing

Genomic DNA was isolated as previously described (Lisch *et al*. 1995). Two micrograms of genomic DNA from the appropriate genotype were digested with restriction enzymes (*Xho*I and *Bam*HI) that cut just outside of the region of interest. Bisulfite conversion was performed using an EpiTect Bisulfite kit (Qiagen). PCR fragments from TIRA were amplified using p1bis2f and TIRAbis2R, and re-amplified using TIRAbis2R and the nested primer TIRAmF6 or p1bis7Fmed (all primer sequences are provided in Supplemental Table 5). In addition, all samples were also amplified using a different set of primers (TIRAMF6 and Autr1R, followed by re-amplification with Autr1R and the nested primers TIRAF1 or p1bis7Fmed). The sequences from each amplification were then compared. This was done to ensure that biases had not been introduced due to the selection of a particular pair of primers. No substantive differences were detected and thus clones from each set of primers were combined. In addition, for many of the plants examined, both fully mature leaves (the third from the top leaf) and immature ears were examined. The results in all cases (different primers or different tissues) were substantially equivalent. An example of results obtained using two sets of primers on both leaves and immature ears from one plant is portrayed in Figure S4. To ensure that duplicate clones resulting from amplification did not skew the analysis, sequences with zero mismatches were only counted once for each sample. The resulting sequences were analyzed using kismeth (http://katahdin.mssm.edu/kismeth/revpage.pl) (Gruntman *et al*. 2008).

For *Mu1* methylation analysis, a similar strategy was employed, but the initial use of restriction enzymes was not necessary. Following bisulfite conversion, the DNA was amplified using primers Mu1bis1 (located in the *Mu1* element) and either a1mum2bis1 or a1mum2bis2 (located in the *a1-mum2* allele, flanking the *Mu1* insertion).

### Small RNA sequencing

Plants were grown in a greenhouse with a 12 hour light cycle. Young leaf tissue for small RNA samples was obtained from two closely related families segregating for *MuDR* and *Muk*. Each sample class contained at least three pooled individuals, each one of which was genotyped. RNA was independently extracted from two separate sets of individuals on different days, and each set is referred to as an experimental replicate, the results of which are presented separately because of large overall differences in relative abundance of normalized *MuDR*-identical small RNAs. The classes included *MuDR* by itself, *Muk* by itself, *MuDR* with *Muk*, and neither *MuDR* nor *Muk*. The small RNA extraction and enrichment protocol was adapted from Dalmay et. al. (Dalmay *et al*. 2000). Total RNA was extracted using an SDS-based extraction solution and precipitated using ethanol. The pellet was dissolved in water, heated to 65°C for 5 min, and then placed on ice.

Polyethylene glycol (molecular weight 8000; Sigma) was added to a final concentration of 5% and NaCl to a final concentration of 0.5 M. After an hour incubation on ice, the RNA was centrifuged at 10,000*g* for 10 min. Three volumes of ethanol were added to the supernatant, and the RNA was precipitated at –20°C for at least 2 hrs. The low molecular weight RNAs were pelleted by centrifugation for 10 min at 10,000g. Small RNAs were detected in Polyacrylamide gel electrophoresis (PAGE) gel and purified by ZR small-RNA™ PAGE Recovery Kit. Small RNA library was prepared by ScriptMiner™ Small RNA-Seq Library Preparation Kit for Ilumina sequencing.

### Small RNA data analysis

The small RNA sequencing data from different libraries was trimmed and filtered for low quality reads, adapter sequences, and reads matching structural noncoding RNAs (t/r/sn/snoRNAs) collected from Rfam (http://rfam.sanger.ac.uk). The kept reads with a length of 18-26 nt were mapped to *MuDR(p1)* and *Muk*, and their flanking 500 bp upstream and downstream sequences using Bowtie only allowing perfect matches (Langmead *et al*. 2009). Small RNA abundance was normalized to reads per million. The data was viewed using Intergrative Genomics Viewer (Robinson et al. 2011). Small RNAs for the *mop1* and *lbl1* mutants were downloaded from previous studies (Nobuta et al., 2008; Dotto et al., 2014), trimmed and mapped to *MuDR* sequences.

### Statement on reagent and data availability

All small RNA data generated for this work is freely and publicly available. The small RNA sequencing data have been deposited at the National Center for Biotechnology Information Gene Expression Omnibus under accession number GSE103833. Maize lines used for these analyses are also freely available for non-commercial purposes.

---

**Figure S1.**
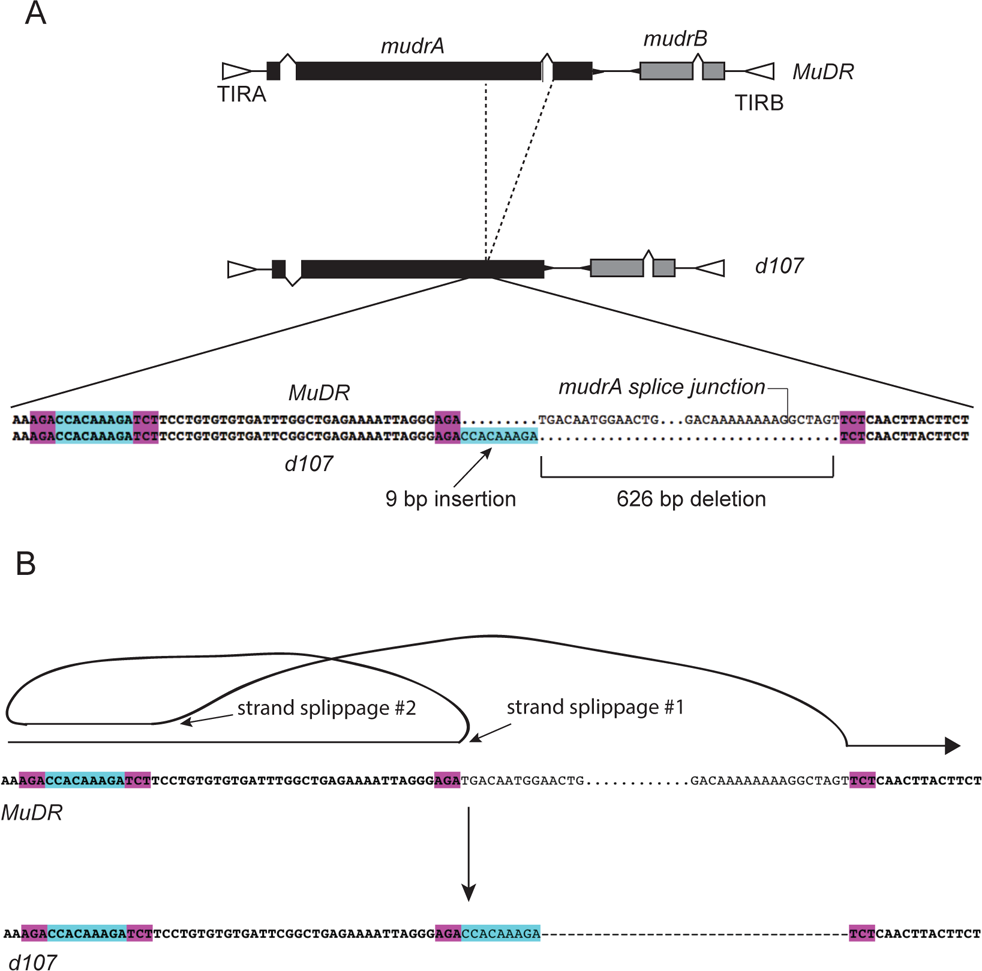
Sequence analysis of the deletion in *d107*. A) Structure of the progenitor (*MuDR*) sequence and that of the deletion derivative. B) Hypothesized mechanism of deletion. *MuDR* replication is thought to involve gap repair using the sister chromatid as a template (Li *et al*. 2008). To generate the deletion at *d107*, we hypothesize that replication proceeded to an AGA triplet, at which point the replicated strand is hypothesized to have switched to a second AGA 47 bp upstream. Replication then continued until it reached a TCT triplet, at which point replication switched to a second TCT triplet 660 bp downstream. The net result was a deletion of 626 bp and the insertion of a short (9 bp) upstream sequence at the end of the deletion.

**Figure S2.**
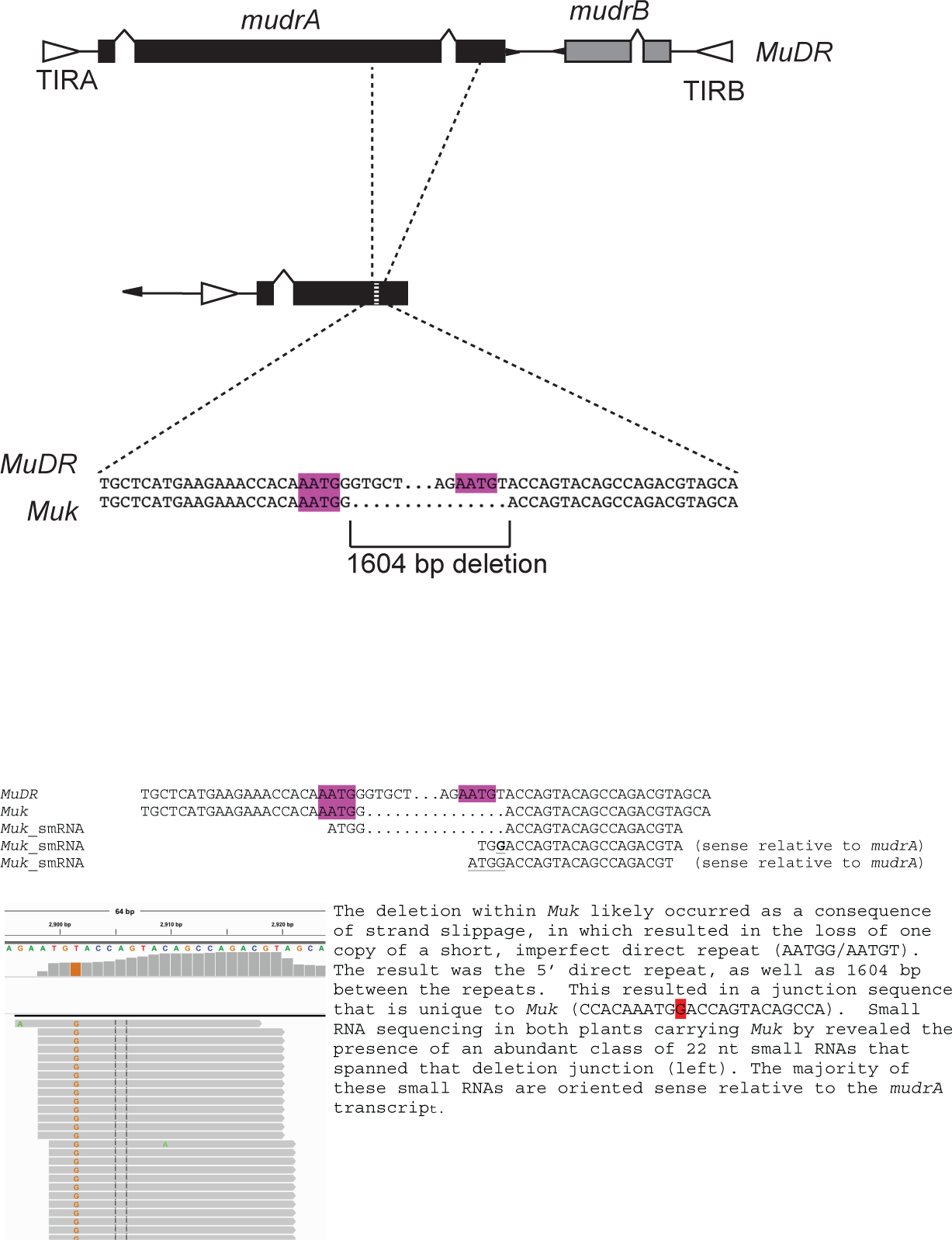
The sequences flanking the deletion within *Muk*, illustrating polymorphisms introduced by that deletion in the small RNA population. As with *d107*, the deletion is likely due to strand slippage at a short direct repeat. The result is a *Muk*-specific junction sequence that is the source of numerous 22 nt small RNAs.

**Figure S3.**
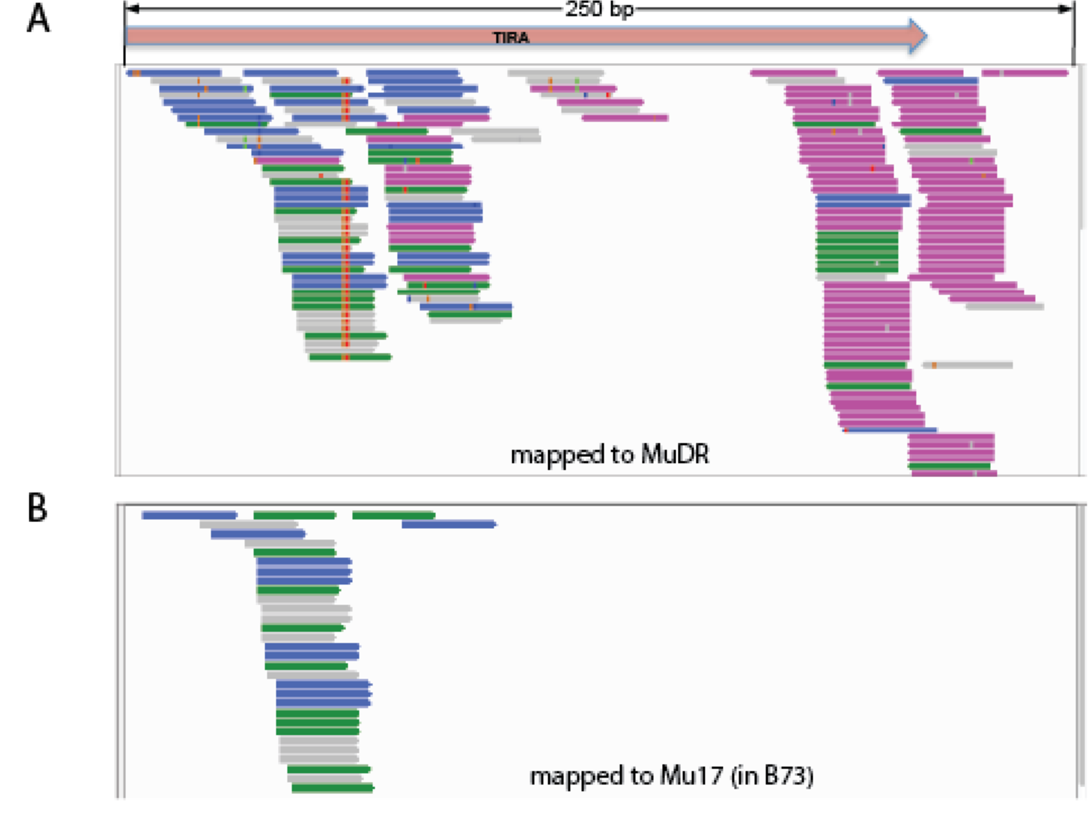
Distribution of small RNAs matched to the 5’ TIR regions of *MuDR(p1)* and *Mu17*. (A) The small RNAs of F1 (p1/-;*Muk/-*) leaf 2 matching the 5’ TIR region of *MuDR(p1)* with two or fewer mismatches. (B) The small RNAs in F1 leaf 2 that perfectly match the 5’ TIR region of Mu17. Green, pink and blue arrows indicate the small RNA length of 21 nt, 22 nt and 24 nt, respectively. The grey arrows represent other sizes of small RNAs.

**Figure S4.**
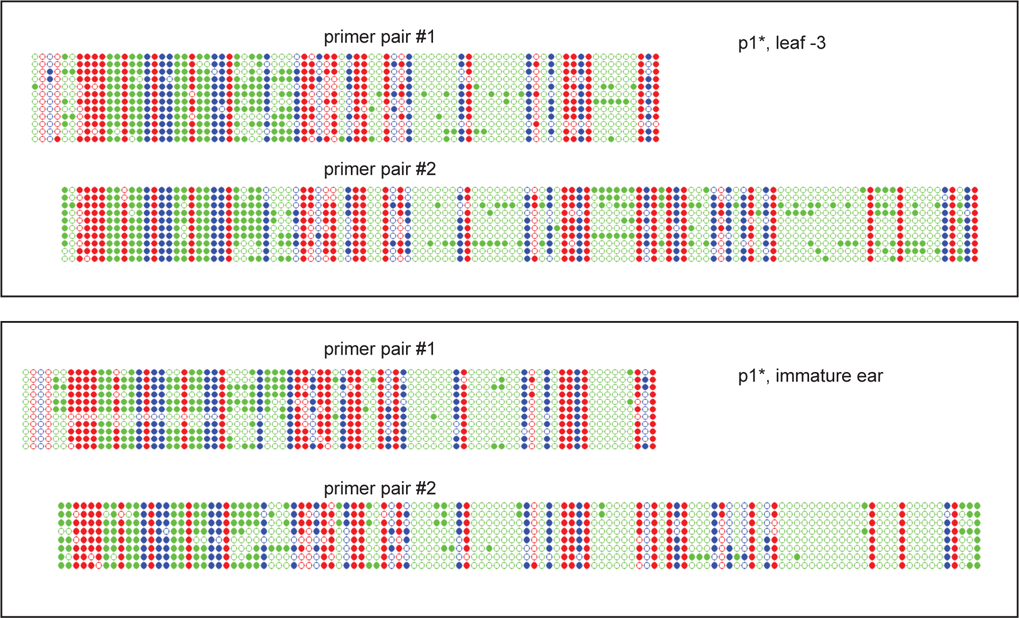
A comparison of the patterns of methylation at TIRA observed in two tissues of the same plant using two different primer pairs. Note that similar results were observed in both tissues regardless of the primer pairs used.

**Supplemental Table 1.**
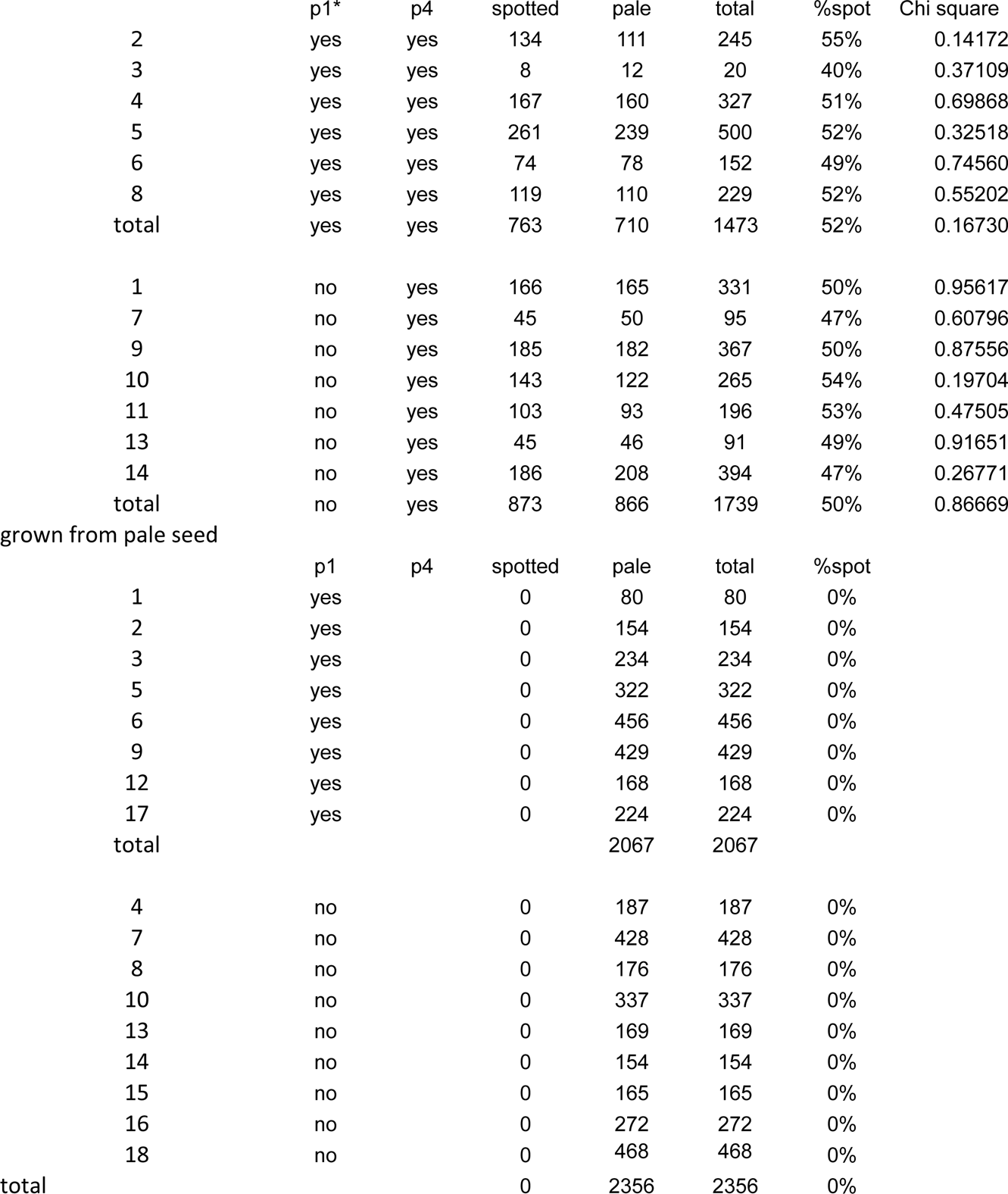
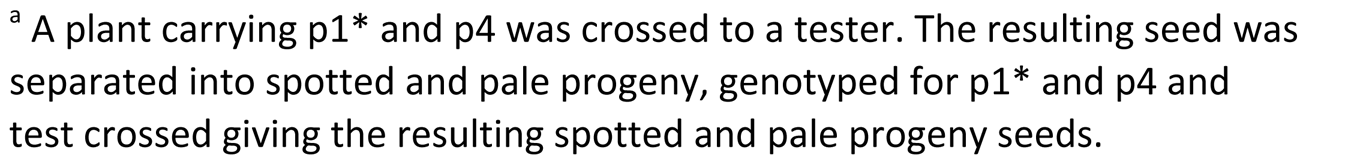
Genetic interaction between a silenced and an active *MuDR* element. grown from spotted seed^a^

|  | p1* | p4 | spotted | pale | total | %spot | Chi square |
| --- | --- | --- | --- | --- | --- | --- | --- |
| 2 | yes | yes | 134 | 111 | 245 | 55% | 0.14172 |
| 3 | yes | yes | 8 | 12 | 20 | 40% | 0.37109 |
| 4 | yes | yes | 167 | 160 | 327 | 51% | 0.69868 |
| 5 | yes | yes | 261 | 239 | 500 | 52% | 0.32518 |
| 6 | yes | yes | 74 | 78 | 152 | 49% | 0.74560 |
| 8 | yes | yes | 119 | 110 | 229 | 52% | 0.55202 |
| total | yes | yes | 763 | 710 | 1473 | 52% | 0.16730 |
| 1 | no | yes | 166 | 165 | 331 | 50% | 0.95617 |
| 7 | no | yes | 45 | 50 | 95 | 47% | 0.60796 |
| 9 | no | yes | 185 | 182 | 367 | 50% | 0.87556 |
| 10 | no | yes | 143 | 122 | 265 | 54% | 0.19704 |
| 11 | no | yes | 103 | 93 | 196 | 53% | 0.47505 |
| 13 | no | yes | 45 | 46 | 91 | 49% | 0.91651 |
| 14 | no | yes | 186 | 208 | 394 | 47% | 0.26771 |
| total | no | yes | 873 | 866 | 1739 | 50% | 0.86669 |
| grown from pale seed |  |  |  |  |  |  |  |
|  | p1 | p4 | spotted | pale | total | %spot |  |
| 1 | yes |  | 0 | 80 | 80 | 0% |  |
| 2 | yes |  | 0 | 154 | 154 | 0% |  |
| 3 | yes |  | 0 | 234 | 234 | 0% |  |
| 5 | yes |  | 0 | 322 | 322 | 0% |  |
| 6 | yes |  | 0 | 456 | 456 | 0% |  |
| 9 | yes |  | 0 | 429 | 429 | 0% |  |
| 12 | yes |  | 0 | 168 | 168 | 0% |  |
| 17 | yes |  | 0 | 224 | 224 | 0% |  |
| total |  |  |  | 2067 | 2067 |  |  |
| 4 | no |  | 0 | 187 | 187 | 0% |  |
| 7 | no |  | 0 | 428 | 428 | 0% |  |
| 8 | no |  | 0 | 176 | 176 | 0% |  |
| 10 | no |  | 0 | 337 | 337 | 0% |  |
| 13 | no |  | 0 | 169 | 169 | 0% |  |
| 14 | no |  | 0 | 154 | 154 | 0% |  |
| 15 | no |  | 0 | 165 | 165 | 0% |  |
| 16 | no |  | 0 | 272 | 272 | 0% |  |
| 18 | no |  | 0 | 468 | 468 | 0% |  |
| total |  |  | 0 | 2356 | 2356 | 0% |  |

<sup>a</sup> A plant carrying p1\* and p4 was crossed to a tester. The resulting seed was separated into spotted and pale progeny, genotyped for p1\* and p4 and test crossed giving the resulting spotted and pale progeny seeds.

**Supplemental Table 2.**
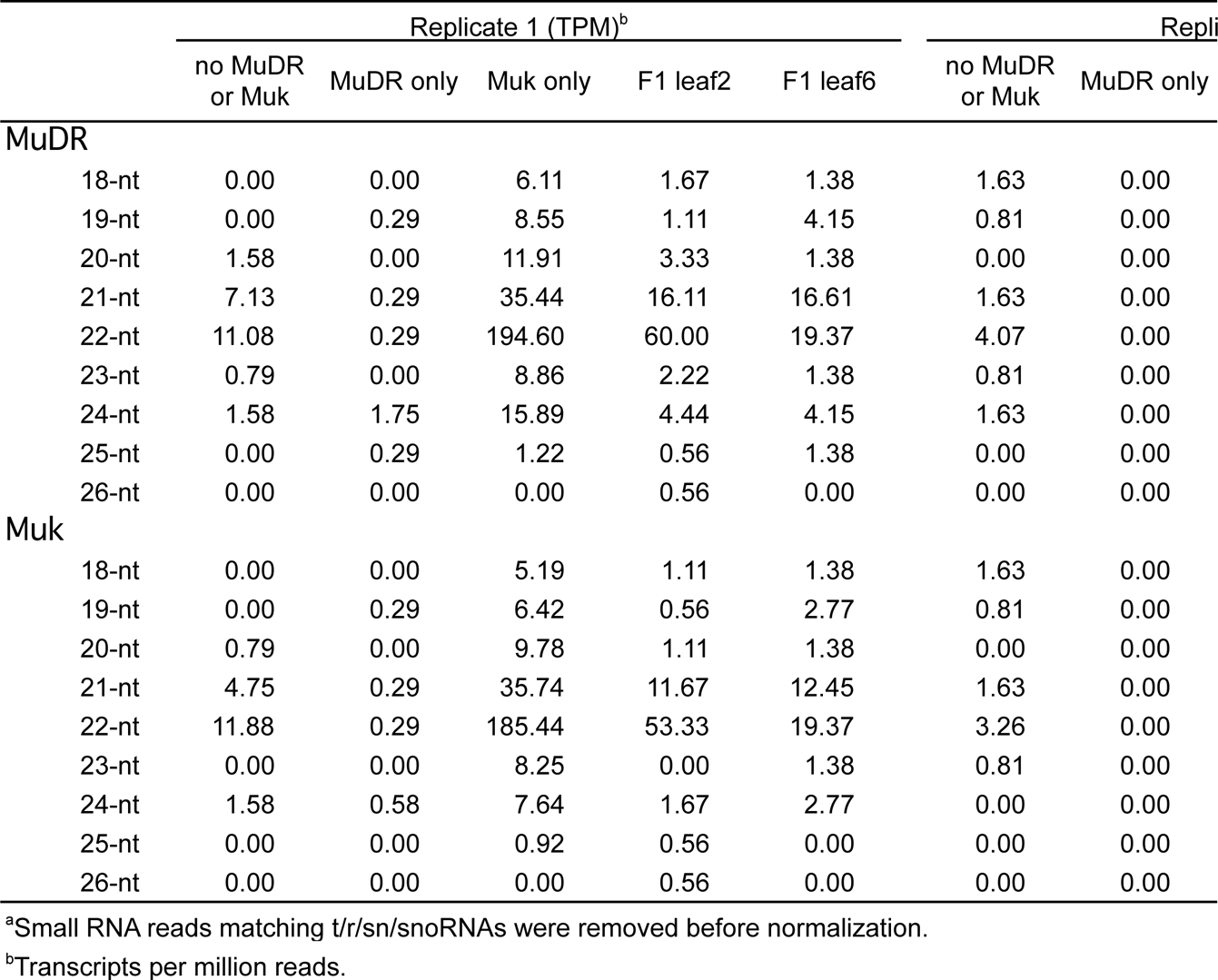
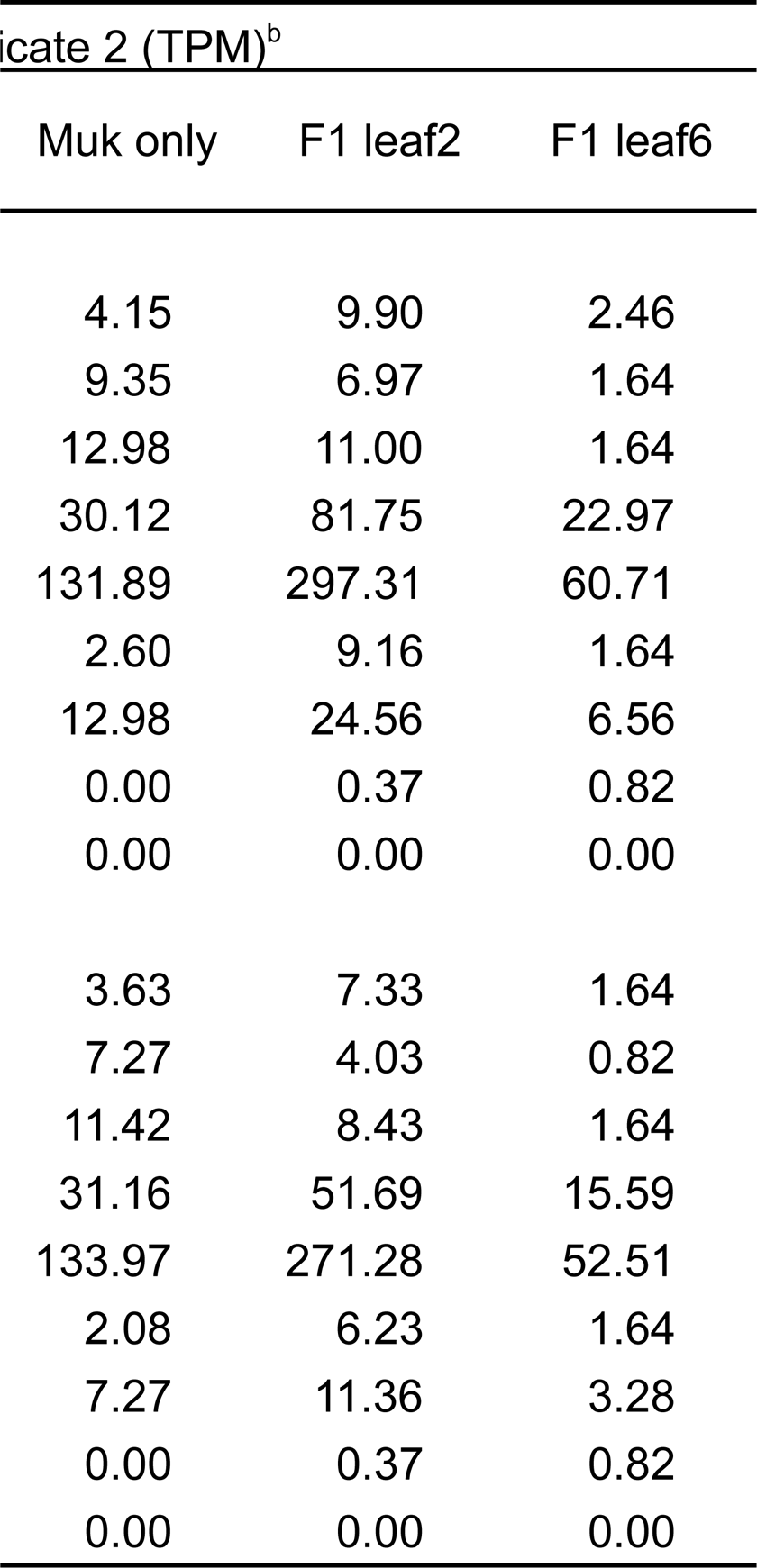
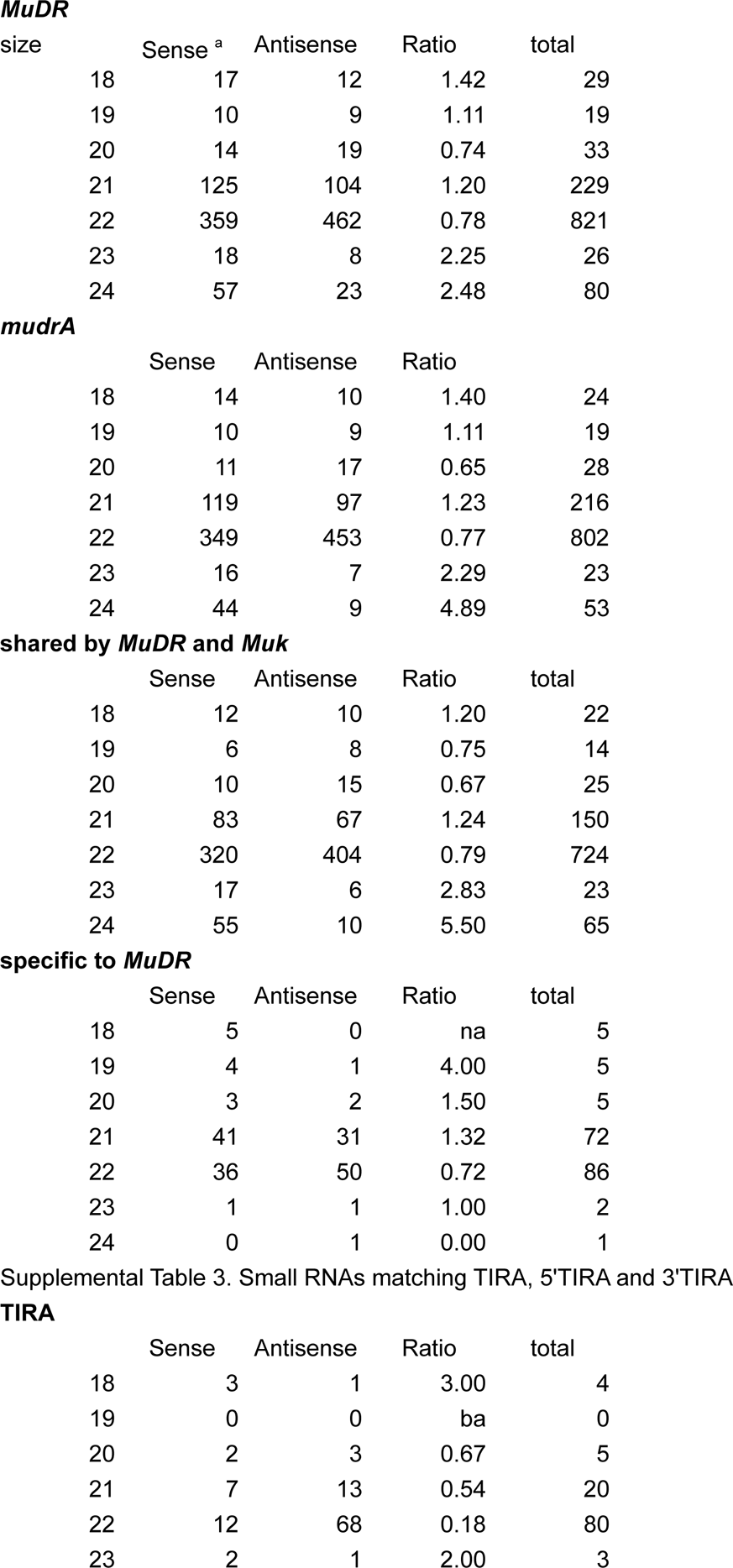
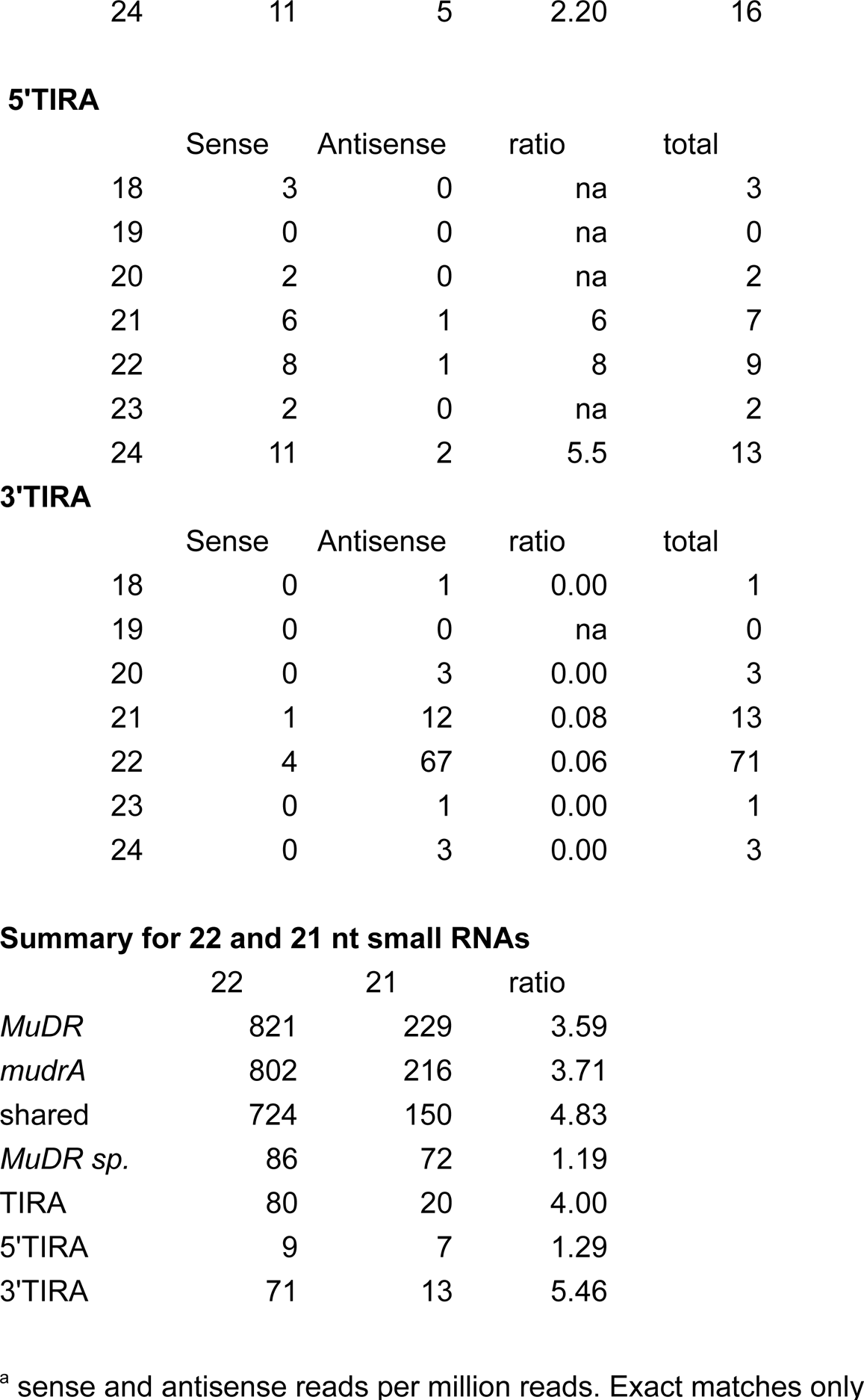
Small RNAs in plants with and without *MuDR* and *Muk*.

**Supplemental Table 4.**
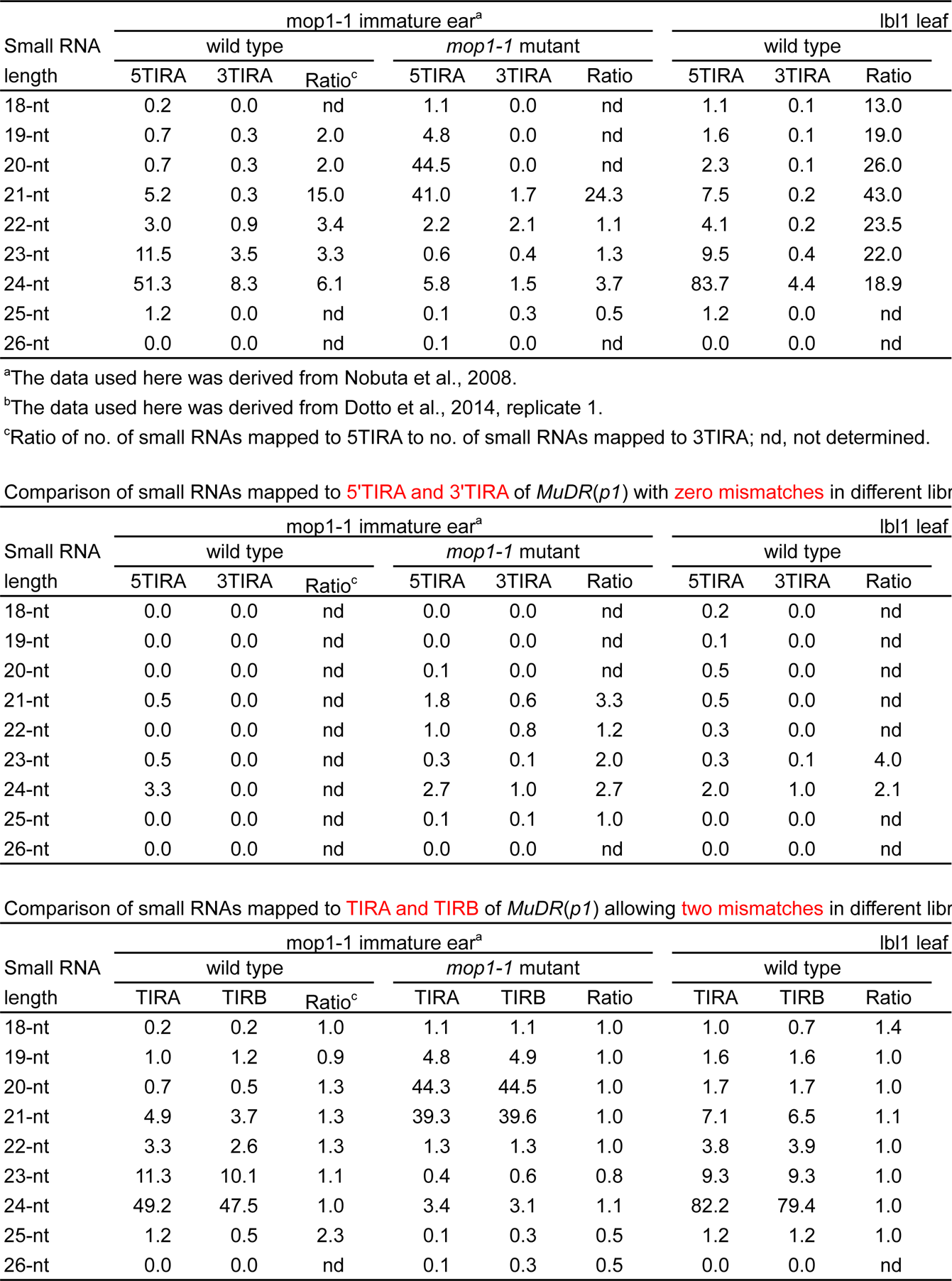
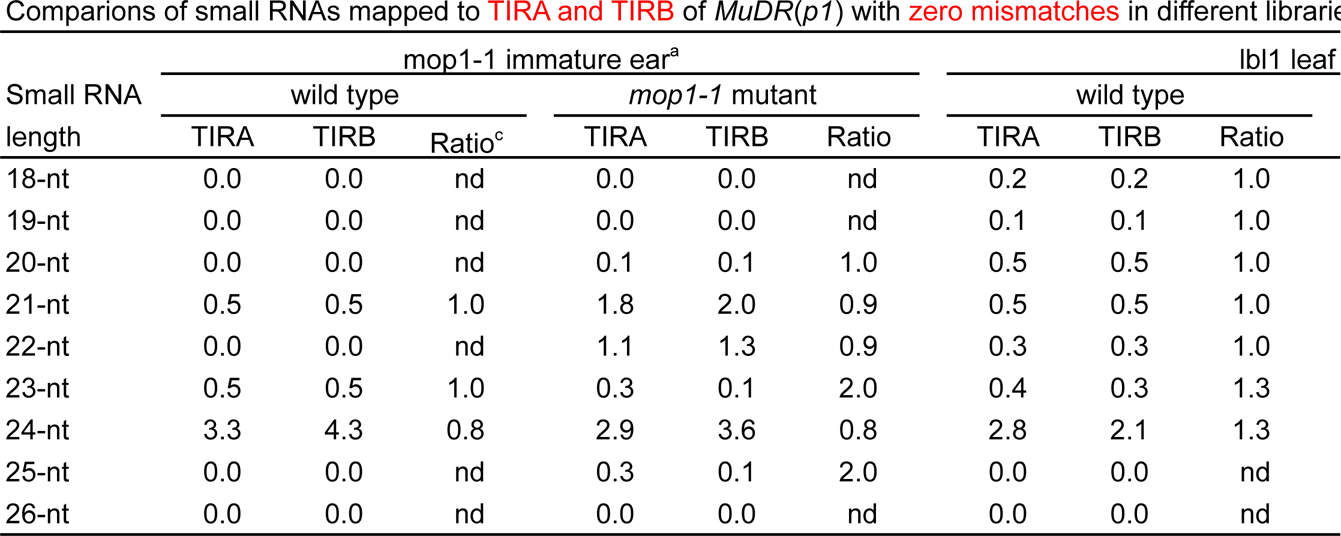
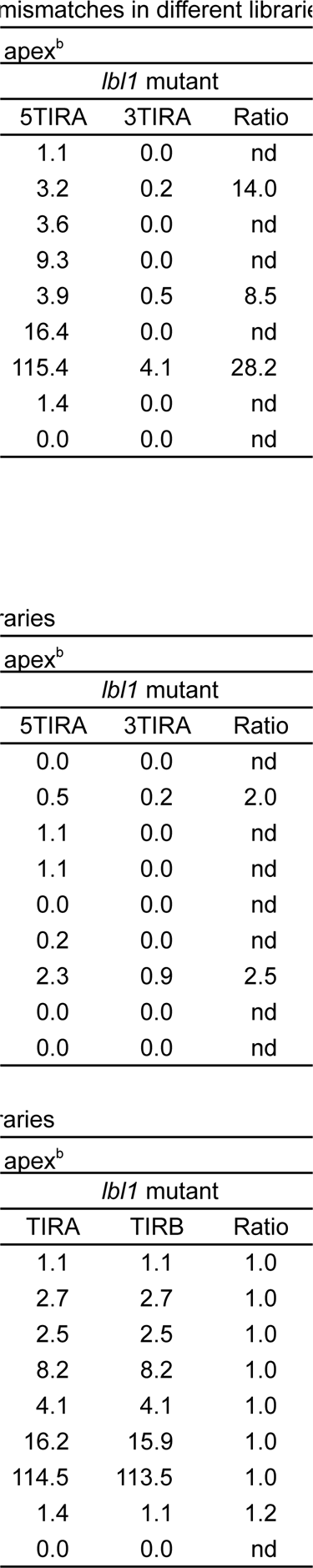
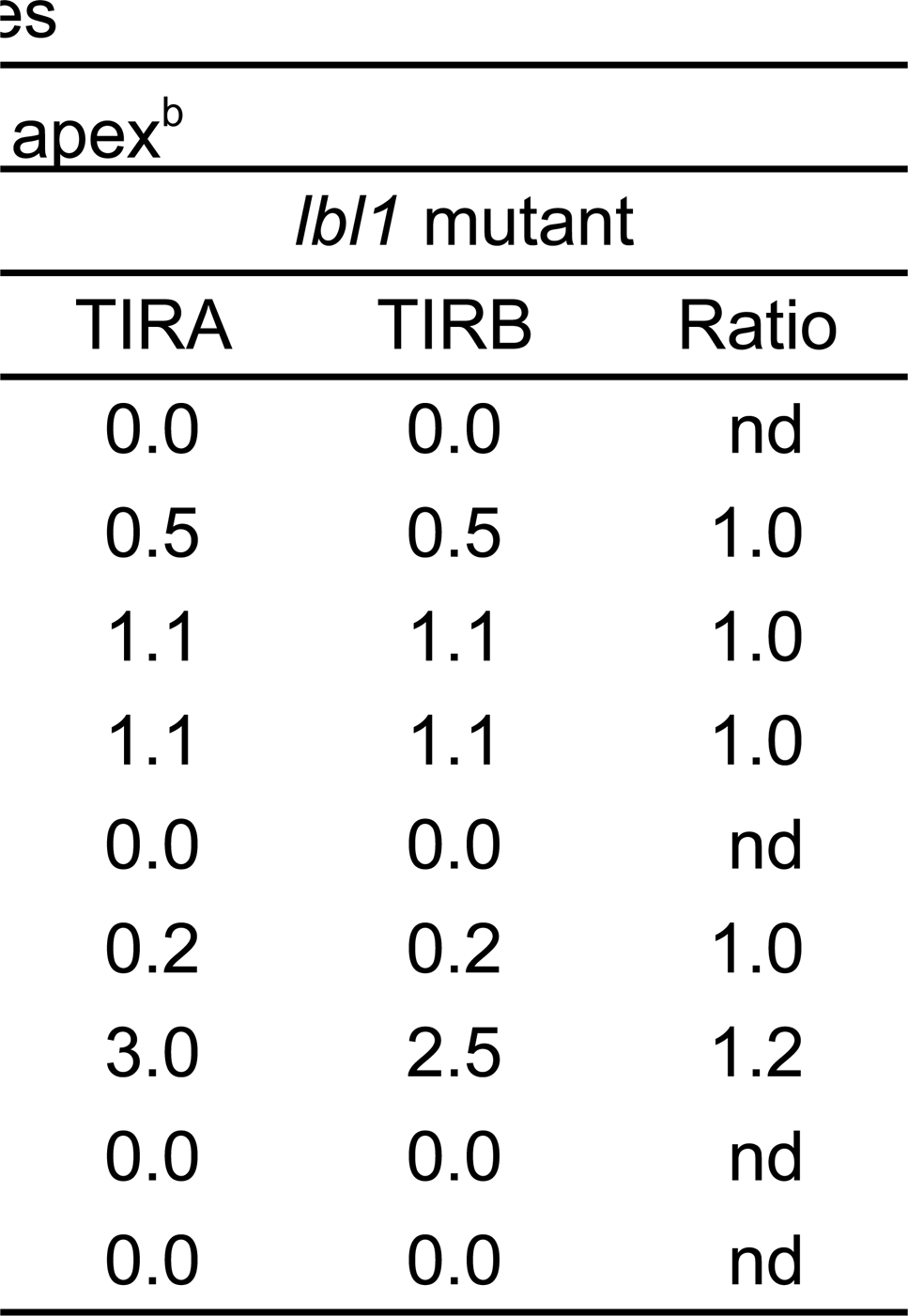
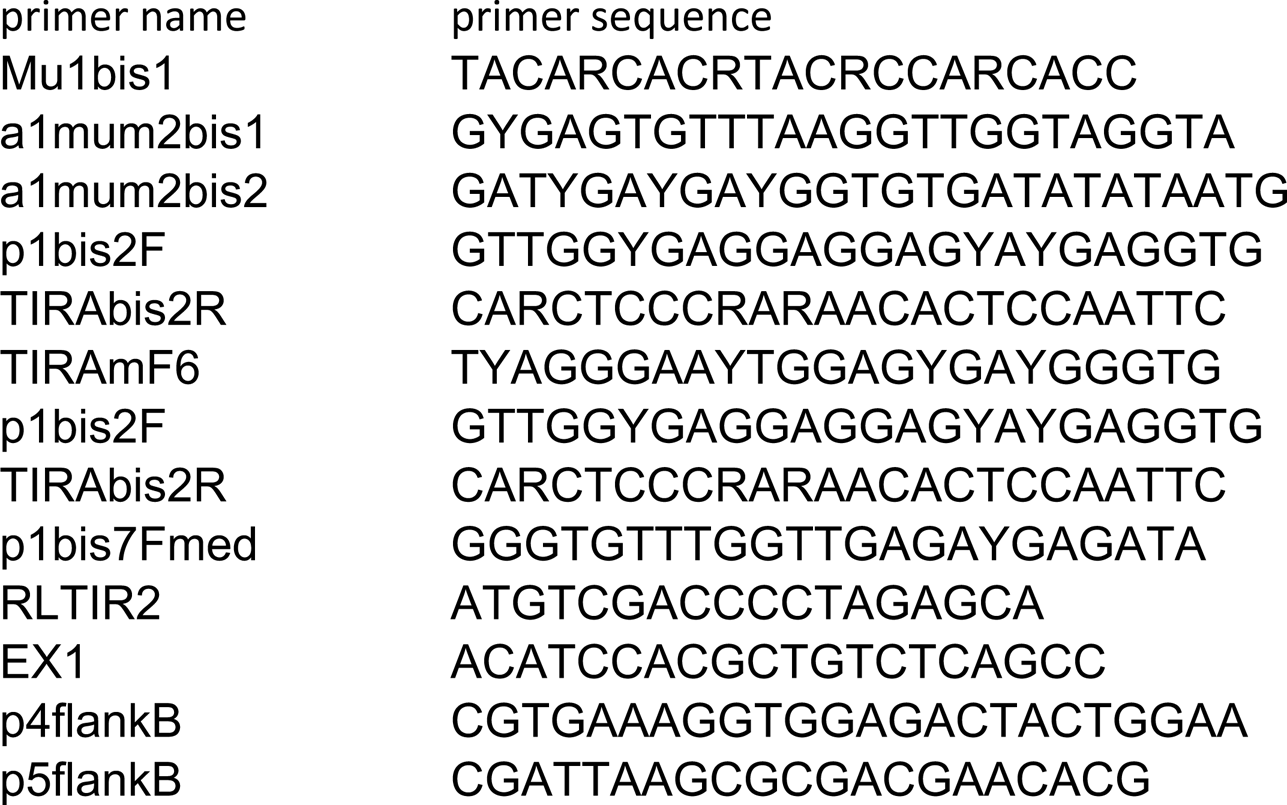
Comparison of small RNAs mapped to 5’TIRA and 3’TIRA of *MuDR*(*p1*) allowing two

## References

1. Alleman, M., L. Sidorenko, K. Mcginnis, V. Seshadri, J. E. Dorweiler et al., 2006 An RNA-dependent RNA polymerase is required for paramutation in maize. Nature 442: 295–298.

2. Banks, J. A., P. Masson and N. Fedoroff, 1988 Molecular mechanisms in the developmental regulation of the maize Suppressor-mutator transposable element. Genes Dev 2: 1364–1380.

3. Barkan, A., and R. A. Martienssen, 1991 Inactivation of maize transposon Mu suppresses a mutant phenotype by activating an outward-reading promoter near the end of Mu1. Proc Natl Acad Sci U S A 88: 3502–3506.

4. Benito, M. I., and V. Walbot, 1997 Characterization of the maize Mutator transposable element MURA transposase as a DNA-binding protein. Mol Cell Biol 17: 5165–5175.

5. Bennetzen, J. L., 1996 The Mutator transposable element system of maize. Curr Top Microbiol Immunol 204: 195–229.

6. Bucher, E., J. Reinders and M. Mirouze, 2012 Epigenetic control of transposon transcription and mobility in Arabidopsis. Curr Opin Plant Biol 15: 503–510.

7. Chandler, V. L., and V. Walbot, 1986 DNA modification of a maize transposable element correlates with loss of activity. Proc Natl Acad Sci U S A 83: 1767–1771.

8. Chomet, P., D. Lisch, K. J. Hardeman, V. L. Chandler and M. Freeling, 1991 Identification of a regulatory transposon that controls the Mutator transposable element system in maize. Genetics 129: 261–270.

9. Creasey, K. M., J. Zhai, F. Borges, F. Van Ex, M. Regulski et al., 2014 miRNAs trigger widespread epigenetically activated siRNAs from transposons in Arabidopsis. Nature 508: 411–415.

10. Cuerda-Gil, D., and R. K. Slotkin, 2016 Non-canonical RNA-directed DNA methylation. Nat Plants 2: 16163.

11. Cui, H., and N. V. Fedoroff, 2002 Inducible DNA demethylation mediated by the maize Suppressor-mutator transposon-encoded TnpA protein. Plant Cell 14: 2883–2899.

12. Dalmay, T., A. Hamilton, E. Mueller and D. C. Baulcombe, 2000 Potato virus Xamplicons in arabidopsis mediate genetic and epigenetic gene silencing. Plant Cell 12: 369–379.

13. Dotto, M. C., K. A. Petsch, M. J. Aukerman, M. Beatty, M. Hammell et al., 2014 Genome-wide analysis of leafbladeless1-regulated and phased small RNAs underscores the importance of the TAS3 ta-siRNA pathway to maize development. PLoS Genet 10: e1004826.

14. Fultz, D., S. G. Choudury and R. K. Slotkin, 2015 Silencing of active transposable elements in plants. Curr Opin Plant Biol 27: 67–76.

15. Fultz, D., and R. K. Slotkin, 2017 Exogenous Transposable Elements Circumvent Identity-Based Silencing, Permitting the Dissection of Expression-Dependent Silencing. Plant Cell 29: 360–376.

16. Gruntman, E., Y. Qi, R. K. Slotkin, T. Roeder, R. A. Martienssen et al., 2008 Kismeth: analyzer of plant methylation states through bisulfite sequencing. BMC Bioinformatics 9: 371.

17. Hale, C. J., K. F. Erhard, Jr., D. Lisch and J. B. Hollick, 2009 Production and processing of siRNA precursor transcripts from the highly repetitive maize genome. PLoS Genet 5: e1000598.

18. Hashida, S. N., T. Uchiyama, C. Martin, Y. Kishima, Y. Sano et al., 2006 The temperature-dependent change in methylation of the Antirrhinum transposon Tam3 is controlled by the activity of its transposase. Plant Cell 18: 104–118.

19. Hershberger, R. J., M. I. Benito, K. J. Hardeman, C. Warren, V. L. Chandler et al., 1995 Characterization of the major transcripts encoded by the regulatory MuDR transposable element of maize. Genetics 140: 1087–1098.

20. Hirochika, H., K. Sugimoto, Y. Otsuki, H. Tsugawa and M. Kanda, 1996 Retrotransposons of rice involved in mutations induced by tissue culture. Proc Natl Acad Sci U S A 93: 7783–7788.

21. Holoch, D., and D. Moazed, 2015 RNA-mediated epigenetic regulation of gene expression. Nat Rev Genet 16: 71–84.

22. Hsia, A. P., and P. S. Schnable, 1996 DNA sequence analyses support the role of interrupted gap repair in the origin of internal deletions of the maize transposon, MuDR. Genetics 142: 603–618.

23. Ito, H., H. Gaubert, E. Bucher, M. Mirouze, I. Vaillant et al., 2011 An siRNA pathway prevents transgenerational retrotransposition in plants subjected to stress. Nature 472: 115–119.

24. Kato, M., K. Takashima and T. Kakutani, 2004 Epigenetic control of CACTA transposon mobility in Arabidopsis thaliana. Genetics 168: 961–969.

25. Kumakura, N., A. Takeda, Y. Fujioka, H. Motose, R. Takano et al., 2009 SGS3 and RDR6 interact and colocalize in cytoplasmic SGS3/RDR6-bodies. FEBS Lett 583: 1261–1266.

26. Langmead, B., C. Trapnell, M. Pop and S. L. Salzberg, 2009 Ultrafast and memory-efficient alignment of short DNA sequences to the human genome. Genome Biology 10.

27. Law, J. A., and S. E. Jacobsen, 2010 Establishing, maintaining and modifying DNA methylation patterns in plants and animals. Nat Rev Genet 11: 204–220.

28. Li, H., M. Freeling and D. Lisch, 2010 Epigenetic reprogramming during vegetative phase change in maize. Proc Natl Acad Sci U S A 107: 22184–22189.

29. Li, J., T. J. Wen and P. S. Schnable, 2008 Role of RAD51 in the repair of MuDR-induced double-strand breaks in maize (Zea mays L.). Genetics 178: 57–66.

30. Lisch, D., 1995 Genetic and Molecular Characterization of the Mutator System in maize, pp. 408 in Plant and Microbial Biology. Univeristy of California, Berkeley, Berkeley, CA.

31. Lisch, D., 2002 Mutator transposons. Trends Plant Sci 7: 498–504.

32. Lisch, D., 2009 Epigenetic regulation of transposable elements in plants. Annu Rev Plant Biol 60: 43–66.

33. Lisch, D., 2015 Mutator and MULE transposons in Mobile DNA III, edited by P. R. NL Craig, A Lambowitz, M Chandler, M Gellert, S. Sandmeyer. American Society for Microbiology Press, Washington, D.C.

34. Lisch, D., C. C. Carey, J. E. Dorweiler and V. L. Chandler, 2002 A mutation that prevents paramutation in maize also reverses Mutator transposon methylation and silencing. Proc Natl Acad Sci U S A 99: 6130–6135.

35. Lisch, D., P. Chomet and M. Freeling, 1995 Genetic characterization of the Mutator system in maize: behavior and regulation of Mu transposons in a minimal line. Genetics 139: 1777–1796.

36. Lisch, D., L. Girard, M. Donlin and M. Freeling, 1999 Functional analysis of deletion derivatives of the maize transposon MuDR delineates roles for the MURA and MURB proteins. Genetics 151: 331–341.

37. Lisch, D., and N. Jiang, 2008 Mutator and Pack-MULEs, pp. 277-306 in The maize handbook - Volume II: Domestication, Genetics and Genomics of Maize, edited by J. L. Bennetzen and S. Hake. Springer, New York, NY.

38. Mari-Ordonez, A., A. Marchais, M. Etcheverry, A. Martin, V. Colot et al., 2013 Reconstructing de novo silencing of an active plant retrotransposon. Nat Genet 45: 1029–1039.

39. Martinez, G., K. Panda, C. Kohler and R. K. Slotkin, 2016 Silencing in sperm cells is directed by RNA movement from the surrounding nurse cell. Nat Plants 2: 16030.

40. Martinez, G., and R. K. Slotkin, 2012 Developmental relaxation of transposable element silencing in plants: functional or byproduct? Curr Opin Plant Biol 15: 496–502.

41. Matsunaga, W., N. Ohama, N. Tanabe, Y. Masuta, S. Masuda et al., 2015 A small RNA mediated regulation of a stress-activated retrotransposon and the tissue specific transposition during the reproductive period in Arabidopsis. Front Plant Sci 6: 48.

42. Matzke, M. A., and R. A. Mosher, 2014 RNA-directed DNA methylation: an epigenetic pathway of increasing complexity. Nat Rev Genet 15: 394–408.

43. Mccue, A. D., K. Panda, S. Nuthikattu, S. G. Choudury, E. N. Thomas et al., 2015 ARGONAUTE 6 bridges transposable element mRNA-derived siRNAs to the establishment of DNA methylation. EMBO J 34: 20–35.

44. Nobuta, K., C. Lu, R. Shrivastava, M. Pillay, E. De Paoli et al., 2008 Distinct size distribution of endogeneous siRNAs in maize: Evidence from deep sequencing in the mop1-1 mutant. Proc Natl Acad Sci U S A 105: 14958–14963.

45. Nuthikattu, S., A. D. Mccue, K. Panda, D. Fultz, C. Defraia et al., 2013 The initiation of epigenetic silencing of active transposable elements is triggered by RDR6 and 21-22 nucleotide small interfering RNAs. Plant Physiol 162: 116–131.

46. O’reilly, C., N. S. Shepherd, A. Pereira, Z. Schwarz-Sommer, I. Bertram et al., 1985 Molecular cloning of the a1 locus of Zea mays using the transposable elements En and Mu1. EMBO J 4: 877–882.

47. Panda, K., L. Ji, D. A. Neumann, J. Daron, R. J. Schmitz et al., 2016 Full-length autonomous transposable elements are preferentially targeted by expression-dependent forms of RNA-directed DNA methylation. Genome Biol 17: 170.

48. Panda, K., and R. K. Slotkin, 2013 Proposed mechanism for the initiation of transposable element silencing by the RDR6-directed DNA methylation pathway. Plant Signal Behav 8.

49. Robertson, D. S., 1986 Genetic studies on the loss of mu mutator activity in maize. Genetics 113: 765–773.

50. Rudenko, G. N., A. Ono and V. Walbot, 2003 Initiation of silencing of maize MuDR/Mu transposable elements. Plant J 33: 1013–1025.

51. Rudenko, G. N., and V. Walbot, 2001 Expression and post-transcriptional regulation of maize transposable element MuDR and its derivatives. Plant Cell 13: 553–570.

52. Saze, H., and T. Kakutani, 2011 Differentiation of epigenetic modifications between transposons and genes. Curr Opin Plant Biol 14: 81–87.

53. Schwab, R., and O. Voinnet, 2010 RNA silencing amplification in plants: size matters. Proc Natl Acad Sci U S A 107: 14945–14946.

54. Sigman, M. J., and R. K. Slotkin, 2016 The First Rule of Plant Transposable Element Silencing: Location, Location, Location. Plant Cell 28: 304–313.

55. Singh, J., M. Freeling and D. Lisch, 2008 A position effect on the heritability of epigenetic silencing. PLoS Genet 4: e1000216.

56. Skibbe, D. S., J. F. Fernandes and V. Walbot, 2012 Mu killer-Mediated and Spontaneous Silencing of Zea mays Mutator Family Transposable Elements Define Distinctive Paths of Epigenetic Inactivation. Front Plant Sci 3: 212.

57. Slotkin, R. K., 2005 The Heritable silencing of Mutator Transposons by Mu killer, pp. 223 in Plant and Microbial Biology. University of California, Berkeley, Berkeley, CA.

58. Slotkin, R. K., M. Freeling and D. Lisch, 2003 Mu killer causes the heritable inactivation of the Mutator family of transposable elements in Zea mays. Genetics 165: 781–797.

59. Slotkin, R. K., M. Freeling and D. Lisch, 2005 Heritable transposon silencing initiated by a naturally occurring transposon inverted duplication. Nat Genet 37: 641–644.

60. Slotkin, R. K., and R. Martienssen, 2007 Transposable elements and the epigenetic regulation of the genome. Nat Rev Genet 8: 272–285.

61. Slotkin, R. K., M. Vaughn, F. Borges, M. Tanurdzic, J. D. Becker et al., 2009 Epigenetic reprogramming and small RNA silencing of transposable elements in pollen. Cell 136: 461–472.

62. Tan, B. C., Z. Chen, Y. Shen, Y. Zhang, J. Lai et al., 2011 Identification of an active new mutator transposable element in maize. G3 (Bethesda) 1: 293–302.

63. Woodhouse, M. R., M. Freeling and D. Lisch, 2006 Initiation, establishment, and maintenance of heritable MuDR transposon silencing in maize are mediated by distinct factors. PLoS Biol 4: e339.

64. Wu, L., L. Mao and Y. Qi, 2012 Roles of dicer-like and argonaute proteins in TAS-derived small interfering RNA-triggered DNA methylation. Plant Physiol 160: 990–999.

65. Zhao, Z. Y., and V. Sundaresan, 1991 Binding sites for maize nuclear proteins in the terminal inverted repeats of the Mu1 transposable element. Mol Gen Genet 229: 17–26.

